# Modality-specific frequency band activity during neural entrainment to auditory and visual rhythms

**DOI:** 10.1101/2020.11.29.402701

**Authors:** Daniel C. Comstock, Jessica M. Ross, Ramesh Balasubramaniam

## Abstract

Rhythm perception depends on the ability to predict the onset of rhythmic events. Previous studies indicate beta band modulation is involved in predicting the onset of auditory rhythmic events (Snyder & Large, 2005; Fujioka et al., 2009, 2012). We sought to determine if similar processes are recruited for prediction of visual rhythms by investigating whether beta band activity plays a role in a modality dependent manner for rhythm perception. We looked at source-level EEG time-frequency neural correlates of prediction using an omission paradigm with auditory and visual rhythms. By using omissions, we can separate out predictive timing activity from stimulus driven activity. We hypothesized that there would be modality specific markers of rhythm prediction in induced beta band oscillatory activity, characterized primarily by activation in the motor system specific to auditory rhythm processing. Our findings suggest the existence of overlapping networks of predictive beta activity based on common activation in the parietal and right frontal regions, auditory specific predictive beta in bilateral sensorimotor regions, and visually specific predictive beta in midline central, and bilateral temporal/parietal regions. We also found evidence for evoked predictive beta activity in the left sensorimotor region specific to auditory rhythms. These findings implicate modality dependent networks for auditory and visual rhythm perception. The results further suggest that auditory rhythm perception may have left hemispheric specific mechanisms.

## Introduction

Perceiving a rhythm requires making predictions about the temporal onset of rhythmic events. This ability allows us to dance in time with music, play music with others, detect a musical beat, and notice when timing is off the beat. Common measures of rhythm perception are sensorimotor synchronization (SMS) tasks that involve synchronizing one’s movements to rhythmic stimuli. While most humans have little trouble synchronizing to auditory rhythms accurately, synchronizing to visual rhythms can be more variable. SMS to auditory rhythms are more reliable and adaptive (Chen et al., 2002; Repp, 2003; Repp and Penel, 2004; Lorås et al., 2012), compared with visual flashing rhythms (Repp & Su, 2013; Comstock & Balasubramaniam, 2018. However, when synchronizing movements with rhythmically moving visual stimuli such as a bouncing ball, synchronization accuracy improves, yet not to the level of auditory synchronization (Hove et al., 2010; Hove et al., 2013b; Iversen et al., 2015; Gan et al., 2015). The reasons for the disparity in SMS accuracy across auditory and visual modalities are as of yet unclear, and a closer investigation of these mechanisms is required for a complete understanding of neural timing and synchronization processes. The present study aims to explore neurophysiological mechanisms of auditory and visual entrainment, particularly with regard to prediction of rhythmic events.

Previous research has shown there is overlap in the structures involved between visual and auditory rhythm perception, particularly within the premotor cortex, putamen, and cerebellum (Hove et al., 2013a; Araneda et al., 2017). While these areas appear to play a supramodal role in rhythm perception, putamen activation is stronger for auditory rhythms than for visual rhythms, suggesting the auditory system may be more tightly connected to timing networks (Hove et al., 2013a; Araneda et al., 2017). There is also evidence suggesting the visual system has its own in-house rhythm timing mechanisms with sources in the parietal lobes (Jäncke et al., 2000, Jantzen et al. 2005), in MT/V5 (Jantzen et al. 2005) and visual cortex (Zhou et al. 2014, Comstock & Balasubramaniam 2018). Taken together, we interpret this literature as support for modality dependent rhythmic processing mechanisms, although to our knowledge this has not yet been clearly shown with a targeted EEG study.

Beyond modality dependent rhythm processing, it has been suggested timing mechanisms in the brain are task specific (Wiener & Kanai, 2016; Comstock, Hove, & Balasubramaniam, 2018), and may be distinct for aspects of rhythm timing and duration perception (Ross et al., 2018; Grube, Lee et al., 2010; Grube, Cooper et al., 2010). Much of the evidence supporting predictive processing for rhythm comes through measures of neural oscillation within different frequency bands. This oscillatory modulation is believed to indicate communication between different regions of the brain, with lower frequency oscillations involved more in communication between regions that are farther away from each other, and higher frequencies involved more in localized communication (Sarnthein et al., 1998; Von Stein and Sarnthein, 2000). Further, Bastos et al., (2015) have shown in non-human primates that activity in the gamma and theta bands are involved in feedforward, or bottom-up visual processing while the beta band is involved in feedback, or top-down visual processing. Michalareas et al., (2016) have shown similar results in the human visual cortex with gamma involved in bottom up processing and alpha and beta involved in top-down processing. Interestingly, Michalareas et al., (2016) also found that alpha and beta top-down processing affects the ventral and dorsal visual stream areas differently, by shifting dorsal stream activity higher in the functional hierarchy of visual processing, while ventral stream downward. If frequency band activity relates to specific top-down or bottom-up processing networks, then by measuring frequency band activity during different rhythm timing tasks we can find markers of network type involved, supporting different networks for different tasks. Neural oscillation within different frequency bands are therefore a rich source of information for investigating timing networks, astiming information is communicated across brain networks.

Neural mechanisms of auditory rhythm perception rely on strong interactions between motor systems and auditory cortices (Janata et al., 2012; Repp and Su, 2013; Iversen and Balasubramaniam, 2016; Ross et al., 2016a, 2016b), possibly mediated through projections in parietal cortex (Patel and Iversen, 2014; Ross et al., 2018). Communication across these networks could be carried out through frequency band specific oscillatory activity. Activity in the beta band (14 – 30 Hz) is of primary interest as it has been shown to play a role in prediction and timing for auditory rhythms (Snyder & Large, 2005; Fujioka et al., 2009, 2012, 2015), as well as being implicated in the onset of movements (Kilavik et al., 2013).

Snyder & Large (2005) found differentiation between induced and evoked activity in EEG high beta and low gamma bands (20 - 60 Hz), where induced activity is not phase locked to a stimulus onset and evoked activity is phase locked to the stimulus onset. By presenting subjects with a sequence of tones with occasional tones omitted, Snyder and Large found induced activity was similar in tone trials and omitted tone trials, indicating expectation for the tones in the sequence, while evoked activity was greatly reduced when there was no tone. Fujioka et al. (2009) used a similar omission paradigm with MEG and found induced beta from auditory cortices decreased after tone onset and increased in anticipation of the expected tone onset. A later MEG study showed the rate of beta increase in anticipation of tone onset is dependent on the tempo of the stimuli, while beta decrease following tone onset is consistent across multiple tempos (Fujioka et al., 2012). Fujioka et al. (2012) additionally found cortico-cortical coherence that followed the tempo of the rhythms between auditory cortices and sensorimotor cortex, supplementary motor area, inferior-frontal gyrus, and cerebellum.

The role of beta activity in visual rhythm perception is less studied, however, beta band amplitude modulation arising from the motor cortex has also been implicated in visually mediated temporal cues indicating expectation (Saleh et al., 2010). More recently, Varlet et al. (2020), showed cortico-muscular coupling of beta-band activity induced by audio-visual rhythms between EEG recorded over motor areas and EMG recorded from finger muscles pressing down on a force sensor. Significantly, the coupling appeared to be modulated by the tempo of the rhythm and peaked roughly 100 ms prior to each tone in the sequence. Interestingly, the study did not find significant cortico-muscluar coupling in response to separate auditory or separate visual rhythms. While Saleh et al. (2010) and Varlet et al. (2020) suggest involvement of beta band modulation in visual rhythm perception, the role of beta band activity in visual rhythm perception remains unclear.

In order to investigate predictive mechanisms of rhythm perception across modalities, we used EEG to record beta band modulation during auditory and visual rhythms. To separate out the stimulus response activity from activity related to temporal prediction of the stimulus we used an omission paradigm similar to that used by Snyder & Large (2005) and Fujioka et al. (2009). Given that previous studies have indicated involvement of sensorimotor beta in rhythm perception (Fujioka et al., 2012, 2015; Varlet et al., 2020) we investigate both sensory and motor related beta. Because EEG activity smears at the scalp it can be difficult to separate out concurrent sources of activity. We use Independent Components Analysis (ICA) as a blind source separation method in an attempt to distinguish sensory and motor related activity.

Based on the assumption that beta oscillations play a general role in top-down processing, we hypothesized that we would find induced beta power modulation for both auditory and visual modalities following the same pattern seen in Fujioka et al. (2009). Specifically, we hypothesized we would find an induced increase in beta in anticipation of the onset of each rhythmic stimulus event, and also prior to the expected onset of an omitted event (omission onset), followed by a sharp decrease in beta power after event onset, but not after omission onset. Further, we expect that evoked beta power would increase only in response to stimulus onset and not in anticipation of omission onset. Because the motor system has been implicated in both auditory and visual rhythm perception, and evidence of motor related beta for rhythm perception has been seen for auditory rhythms (Fujioka et al., 2012, 2015), and implicated in visual rhythms (Varlet et al., 2020), we expected to find motor related predictive beta activity for both auditory and visual modalities. We also expected to find distinct network activity in predictive beta for visual rhythm perception, specifically greater evidence for predictive beta in the parietal and visual cortices, given evidence of visual timing activity in these regions (Jäncke et al., 2000; Jantzen et al. 2005; Zhou et al. 2014; Comstock & Balasubramaniam 2018)

## Materials and methods

### Participants

18 subjects participated in the experiment (11 female, average age of 23.6 (20 – 34)) with one being rejected after data collection for poor signal to noise ratio. All participants were right-handed and had typical hearing and typical or corrected vision. The experimental protocol was carried out in accordance with the Declaration of Helsinki. This study was approved by the UC Merced Institutional Review Board for research ethics and human subjects, and all participants gave informed consent prior to testing.

### Task

After subjects gave written consent, they were seated and fitted with a 32 electrode EEG cap. Subjects were then tasked with watching isochronous flashing visual rhythms or listening to isochronous auditory rhythms. Both kinds of rhythms had an interonset interval (IOI) of 600 ms, and both had occasional omissions of single tones or single flashes. The rhythms were broken into stimulus trains with each train consisting of 100 tones or flashes with 7 omitted tones or flashes placed randomly within the train. The location of the omitted tones or flashes in the stimulus trains were constrained such that there must be at least 8 tones or flashes between each omission. There were 20 stimulus trains per condition for a total of 140 omissions in each condition. Subjects completed all of the stimulus trains in one modality, followed by all of the stimulus trains in the other modality, in design counterbalanced across subjects. Before the omission conditions, subjects were presented with a condition with no omissions consisting of 140 tones or flashes. The non-omission stimulus trains were of the same modality as the omission stimulus trains that would follow. This design resulted in 140 trials for each of the four conditions (tone non-omission, tone omission, flash non-omission, flash omission).

To ensure that subjects were attending to the rhythms, after each train a shorter sequence of 5 tones or flashes was presented at a slightly slower or faster tempo than the experimental train, and subjects were asked to determine if the shorter rhythm was slower or faster than the preceding rhythm. The number of correct responses and response times were recorded and used to determine if subjects were adequately attending to the stimulus trains.

**Figure 1.**
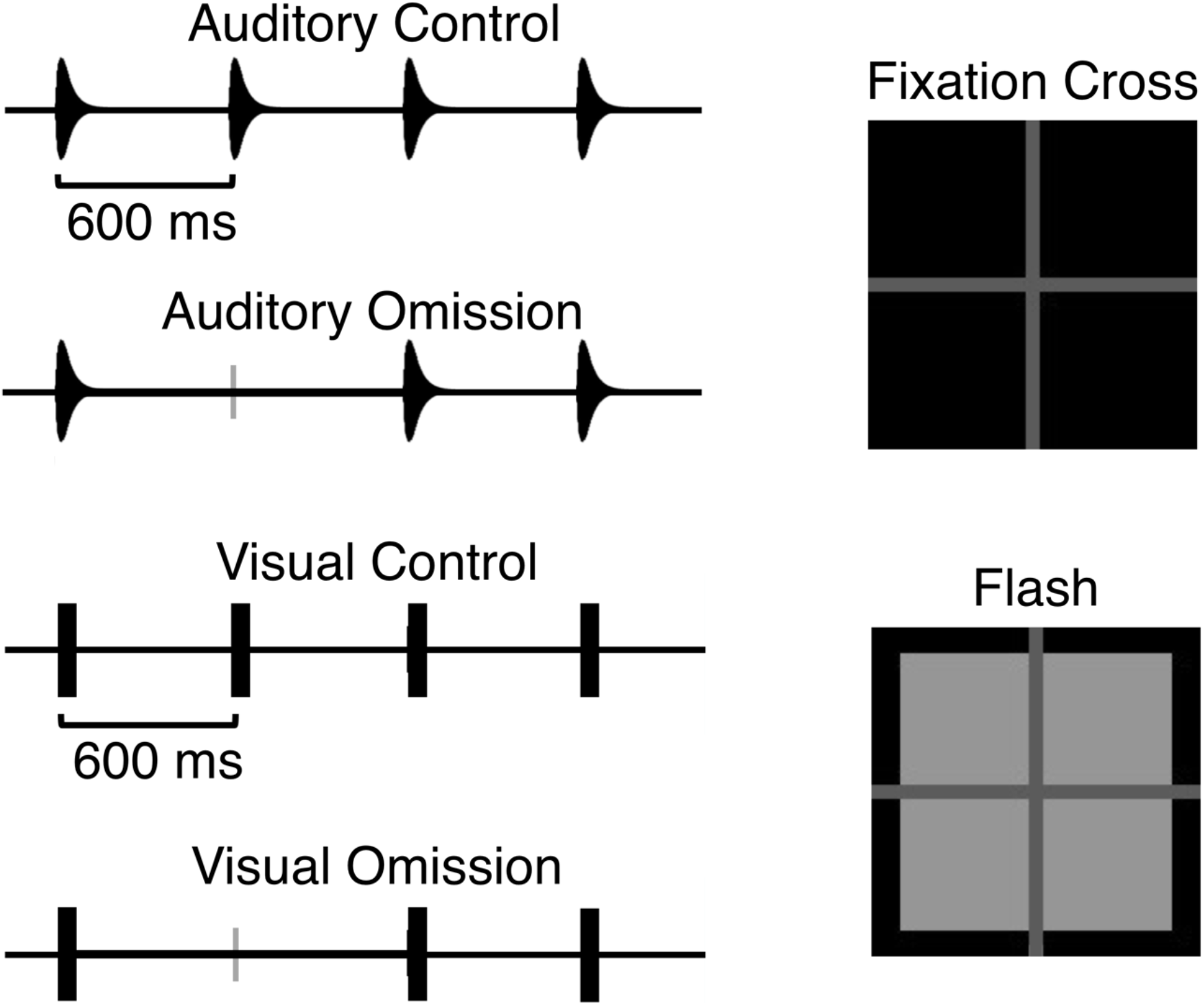
Schematic of control and omission conditions for both auditory and visual metronomes, and depiction of the visual flash metronome stimuli. The fixation cross was always visible for both auditory and visual conditions, even when the flash appeared in the visual condition.

The auditory metronome consisted of 1000 Hz tones lasting 50 ms with a 10 ms rise and 40 ms fall time, generated using Audacity digital audio software. The visual metronome consisted of light grey square flashes 3 cm x 3 cm lasting 50 ms each. In both cases there was a black screen with a dark grey fixation cross in the center of the screen where the lines were approximately 3 mm wide and 4 cm long. The visual flashes always appeared behind the fixation cross so that the cross never disappeared when the flash appeared behind it.

The stimuli were presented using Paradigm experimental stimulus presentation software (Perception Research Systems, 2007) on a 60 Hz monitor, which was approximately 65 cm from the subject’s head. Subjects responded to any prompts using a keyboard placed on a desk in front of the chair they were seated in.

### EEG data acquisition and processing

EEG was continuously recorded using an ANT-Neuro 32 channel amplifier with the ANT-Neuro 32 electrode Waveguard cap. The electrodes were situated according to the 10-20 International system and EEG was recorded with a sampling rate of 1024 Hz. The data were then processed using the EEGLAB v14.1.1 toolbox (Delorme and Makeig, 2004) within Matlab 2019a. Channel locations were added using the standard location montage for the Wavegaurd cap. EEG data were first pruned by hand to remove sections between stimulus train blocks. This was done to remove any break periods between trains. Following pruning, the data were down-sampled to 256 Hz and then a high pass filter with a 2 Hz passband edge and 6 dB cutoff at 1 Hz was applied. A lowpass filter with a 50 Hz passband edge and 6 dB cutoff at 56.25 Hz was applied to remove 60 Hz line noise. Bad channels were rejected that had activity with lower than 0.8 correlation with their surrounding channels, and the rejected channels were then interpolated using spherical interpolation. We then removed single channel artifacts using artifact source reconstruction (ASR) which has been shown to effectively remove large-amplitude or transient artifacts in the data (Mullen et al., 2015; Chang et al., 2018). ASR was performed using a conservative burst criterion parameter of 50 standard deviations. After ASR was run we then rereferenced the data to average. In order to separate out non-brain artifacts and for the source level analysis we ran Independent Components Analysis (ICA) using the AMICA ICA algorithm (Palmer et al., 2012). Dipole source localization was performed on the resulting components using the MNI head model, and 2 dipoles were fit where appropriate instead of 1 using the FitTwoDipoles plug in (Piazza et al., 2016). ICA components were checked to find eye blink and cardiac components, which were marked for later rejection. The remaining independent components were used for source analysis

We then segmented the continuous data into 4 long epochs for the experimental conditions: Non-omission visual flashes, non-omission auditory tones, visual omissions, and auditory omissions. The non-omission conditions came from the non-omission stimulus train block that preceded the omission block. Each condition was epoched from −1.67 seconds prior to each tone/flash to 1.67 seconds following the tone/flash. Epoch length was determined by calculating the window size needed for the later time/frequency calculations so the resulting time/frequency data would span +/− 1.5 seconds from the tone or flash onset of interest. The omission groups were epoched in the same way in relation to omission events. Following epoching, epochs were checked for blinks that occurred during either event onset (for the non-omission conditions) or expected onset (for the omission conditions) as defined as a 50 uV or larger spike in frontal electrodes within +/− 100 ms of onset or expected onset. After epochs with eye blinks at event onset, or expected onset, were rejected, eye blink components determined by AMICA marked earlier were then rejected. Remaining epochs with amplitude spikes greater than +/− 500 uV were then rejected. Finally, epochs that were deemed improbable were rejected by computing the probability distribution of values across the epochs for individual channels and across all channels. Any epoch that contains data values greater 6 standard deviations for the channel or 2 standard deviations for all electrodes was rejected. One subject was rejected due to having over 50 % of their total epochs being rejected. For the remaining 17 subjects there were 140 possible epochs per condition per subject for the 4 conditions: Visual Non-omission (M = 123.24, max = 136, min = 96, MAD = 13.27), Visual Omission (M= 116.59, max = 132, min = 74, MAD = 18.19), Auditory Non-omission (M = 118.29, max = 136, min = 92, MAD = 14.24), Auditory Omission (M = 109.06, max = 129, min = 66, MAD = 20.12).

EEG activity measured at the electrode level is smeared across the scalp making it difficult to separate out signals from different sources. Because we are interested in time sensitive neural activity from both sensory and motor areas that occur simultaneously, we focus our analysis on the source level components. To compare independent components across subjects, we performed a cluster analysis using k-means clustering based on the component dipole locations and scalp topographies. To ensure non-brain sources were excluded from clustering, only components with dipoles located within the head and with a residual variance of less than 15% were used resulting in a total of 289 total brain components across 17 subjects. To determine the appropriate number of clusters, we applied three measures for cluster number optimization (Calinski-Harabasz, Silhouette, and Davies-Bouldin) for between 5 and 30 clusters. The Calinski-Harabasz and Silhouette methods indicated the optimal number of clusters was 9 while the Davies-Bouldin method indicated an optimum number of 13. We used 9 clusters to maximize the number of unique subjects per cluster, plus 1 outlier cluster with components with positions of more than 3 standard deviations from any of the cluster centers. In addition, the parent cluster consisted of all 289 components. The 9 clusters (figure 2, table 1) averaged 31.78 components per cluster with a standard deviation of 7.1, which were made up from 15.78 subjects on average, standard deviation 0.97. The outlier cluster consisted of three components from 2 subjects.

**Table 1.** Information containing the component make up of the 9 clusters and outlier cluster. Although the corresponding Brodmann area for each cluster is determined based on the average talairach coordinates of the component dipoles, the dipole locations for the individual components for each cluster are not all contained within the indicated Brodmann area. Individual dipoles for each component are shown in figure 2

| Cluster | Subjects | Components | Mean R.V. % | Mean Tal Coordinates | Corresponding Brodmann Area of Mean coordinates |
| --- | --- | --- | --- | --- | --- |
| 1 - Left Frontal | 15 | 20 | 5.88 % | X: -31 Y: 45 Z: 14 | Left Area 10 |
| 2 - Left sensorimotor | 16 | 36 | 5.86% | X: -53 Y: -12 Z: 19 | Left BA 1 / 4 (Primary Sensory / Primary Motor) |
| 3 - Midline Central | 17 | 35 | 6.54 % | X: -12 Y: -12 Z: 51 | Left BA 6 |
| 4 - Right Sensorimotor | 16 | 34 | 5.21% | X: 49 Y: -9 Z: 30 | Right BA 4 (Primary Motor) |
| 5 - Right Frontal | 17 | 39 | 7.07% | X: 18 Y: 32 Z: 20 | Right BA 8 / 9 |
| 6 - Left Temporal / Parietal | 15 | 28 | 4.49% | X: -42 Y: -58 Z: 13 | Left BA 39 (Angular Gyrus) |
| 7 - Occipital | 14 | 24 | 4.52% | X: -4 Y: -87 Z: -5 | Left BA 18 (Visual Assoc) |
| 8 - Parietal | 16 | 42 | 4.42% | X: 10 Y: -58 Z: 45 | Right BA 7 |
| 9 - Right Temporal / Parietal | 16 | 29 | 3.42% | X: 41 Y: -61 Z: 7 | Right BA 19 / 37 (Peristriate Area / Fusiform Gyrus) |
| 10 - (Outlier) | 2 | 3 | 7.03% | X: 10 Y: -31 Z: -19 | Null |

**Figure 2.**
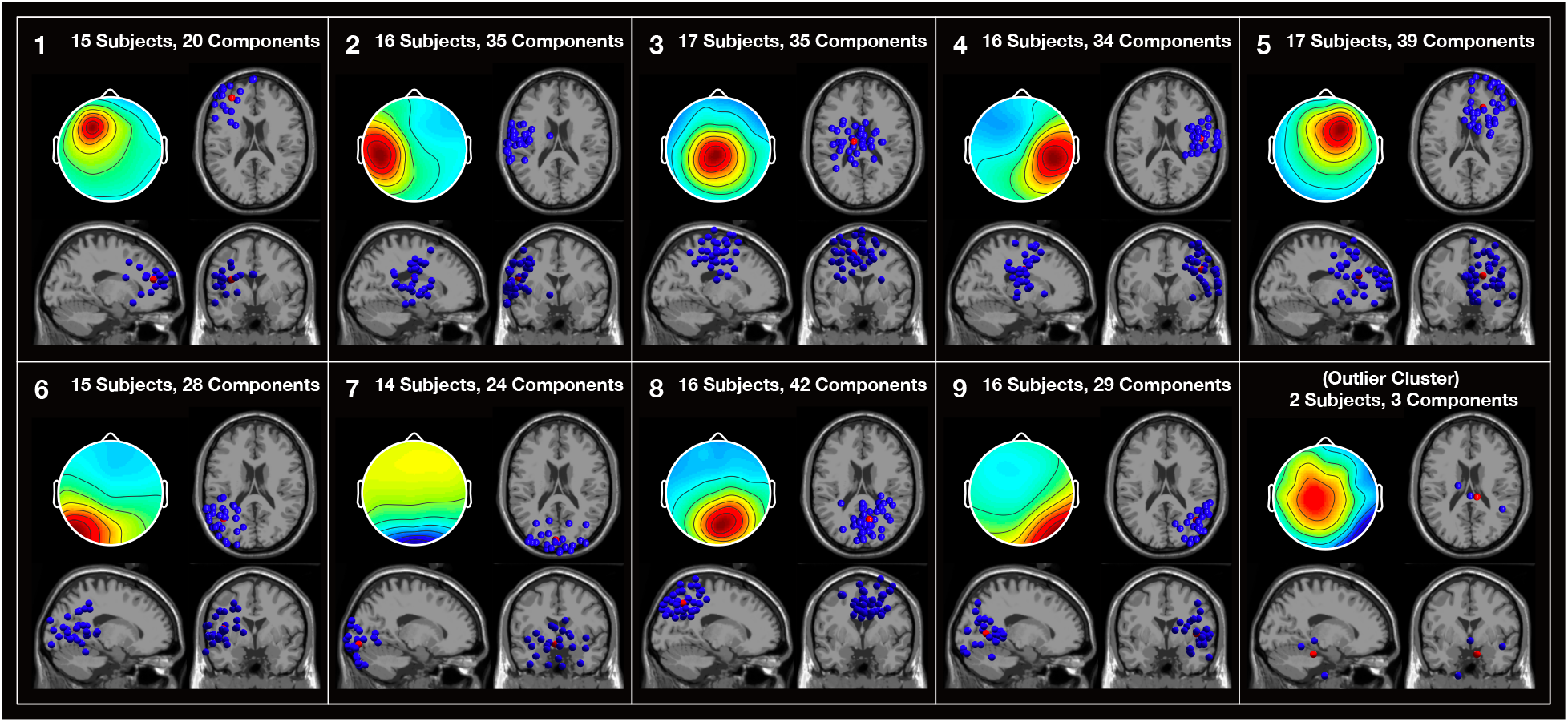
Scalp topography and dipole locations of components for the 9 clusters and the outlier cluster. Scalp topography includes activity from all four conditions. Blue dots indicate individual component dipole locations. Red dots indicate the average position.

### Time frequency analysis

Time frequency analysis was completed for each subject at each channel and for each component used in the clustering analysis. The resulting time frequency representations were then averaged across subjects for the individual channels in each condition, and averaged across the components for each cluster for each condition. Both induced and evoked time frequency representations were calculated to determine the different roles they may play during the rhythm perception task. Induced activity was calculated for each trial by first removing the mean of activity (ERP) from each trial so only non-phase locked activity remains, and then averaging the resulting time frequency computations across trials. Evoked activity was calculated on the mean of the activity (ERP) to focus on the phase locked activity. Both induced and evoked activity were calculated using the same parameters. The time frequency calculations were computed using 85 linear spaced Morlet wavelets between 8 and 50 Hz with a fixed window size of 300 ms resulting in 2.4 cycles at 8 Hz and scaling up to 15 cycles at 50 Hz. The 300 ms window size was chosen to ensure the time frequency representation from each individual stimulus was not contaminated by either surrounding stimuli, which were 600 ms apart. The convolution used the minimum step size for the sample rate of 256 Hz resulting in 772 evenly spaced steps with a step length of 3.9 ms. A relative to the mean baseline was used with the baseline computed separately for each condition. The baseline period for each condition was taken from −1200 to −600 ms of the stimulus onset and therefore consisted of one complete 600 ms stimulus cycle for both the omission and control conditions. These computations were used to determine the Event Related Spectral Perturbation (ERSP) values in terms of dB, such that the ERSP plots show shift in power from baseline at each time point. Beta activity was extracted from the ERSP values by averaging the power at each step from between 14 and 30 Hz.

## Results

### Attention task behavioral results

To assess if attention was maintained evenly between the two modalities, we analyzed the behavioral data from the attention task for the two omission conditions. Both auditory (94.72%) and visual (88.61%) conditions showed a correct response rate well above chance. To assess the differences between the auditory and visual conditions, the number of correct responses and response times were assessed using paired t-tests. There was a significant difference in number of correct attention trials between the auditory (M = 18.94, SD = 1.09) and visual (M = 17.65, SD = 1.69) conditions; t(16) = −2.72, p = 0.015 which we ascribe to the visual rhythm task being more difficult than the auditory rhythm task. There was no significant difference in response time measured in ms between auditory (M = 1405.04, SD = 572.09) and visual (M = 1495.99, SD = 585.94) conditions; t(16) = 0.66, p = 0.52.

### Event Related Spectral Perturbations

To determine if ERSP power was being significantly modulated by the stimuli and omissions, permutation statistics comparing ERSP power values to baseline values using unpaired t-tests with 2000 permutations testing for significance were performed. FDR correction was used to correct for multiple comparisons with alpha values being the computed p-value for each time-frequency point using a parametric FDR algorithm. In the ERSP plots (Figures 3–7), areas within the black lines in the induced ERSP plots correspond to p <0.01 values. Because the amount of power modulation was much greater in the evoked ERSP power, areas within the black lines in the evoked ERSP plots correspond to p < 0.001 values.

**Figure 3.**
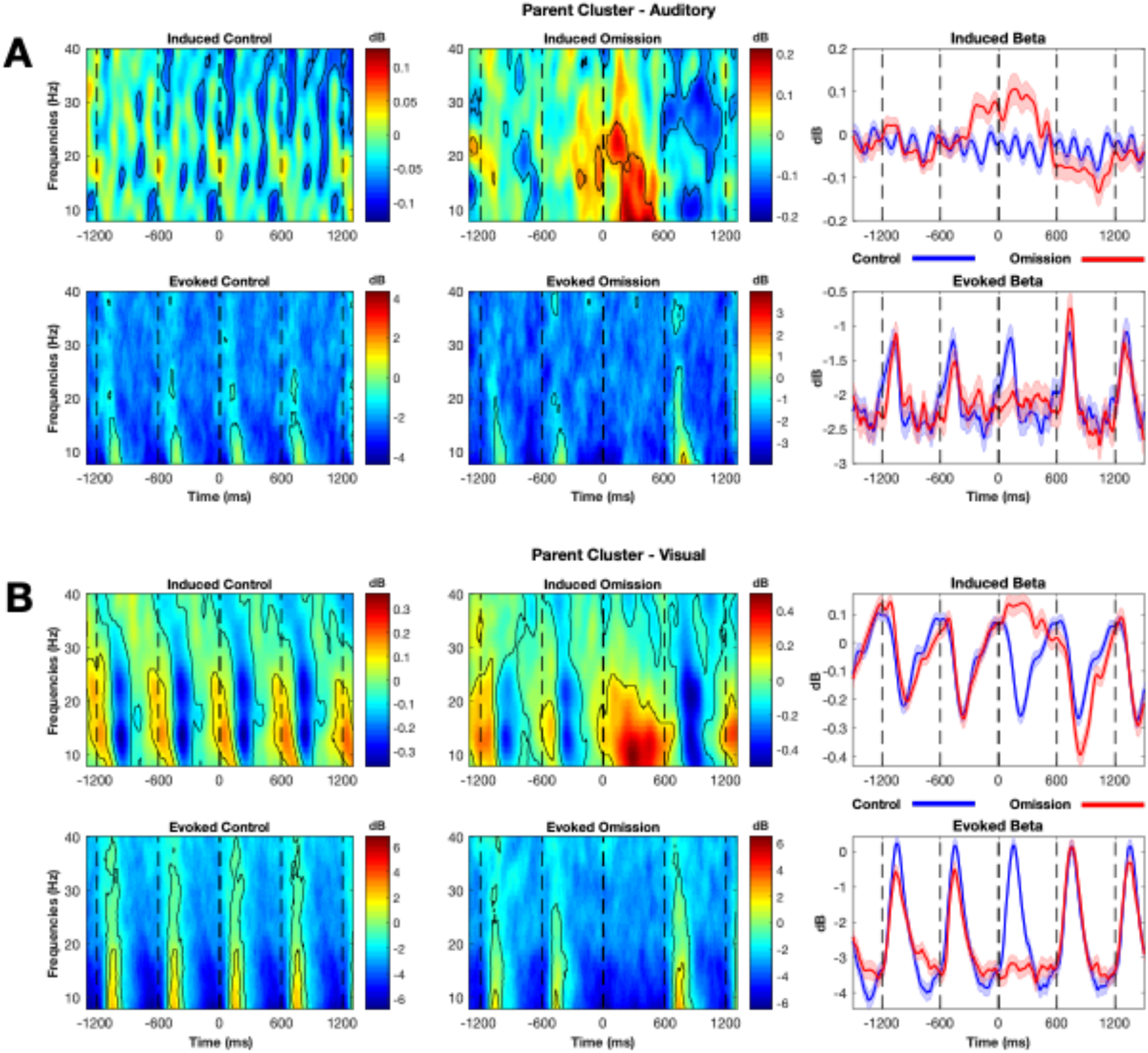
Induced and evoked ERSP and beta band time frequency representations for both Auditory (A) and Visual (B) conditions for the parent cluster which contains all components from all subjects. Significant time frequency values in the ERSP plot are outlined in black Both auditory and visual Evoked beta power appears to respond only to stimulus onset and not to the omission onset. Induced beta power increases prior to visual stimulus onset and prior to the omitted onset. Shaded bars in beta plots indicate standard error.

Looking at the parent cluster containing all components, we find increased evoked power following both visual and auditory stimulus onset, but not in response to visual or auditory omission onsets (Figure 3a & 3b). Induced activity from the visual condition in the parent cluster increases significantly and peaks roughly at stimulus onset, but also increases at omission onset, particularly in the low beta range (Figure 3b). This pattern is also seen in the posterior clusters for visual activity (Figures 4 & 5). Auditory ERSP power modulation is less pronounced compared to visual, and although evoked activity in the parent cluster increases in response to auditory stimuli in the beta range, induced power does not significantly increase at stimulus onset for auditory beta as it does for visual beta (Figure 3a). Auditory evoked power in the beta range appears to increase after both stimulus onset and omission onset in the left and right sensorimotor area clusters (Figures 5A & 6 A).

**Figure 4.**
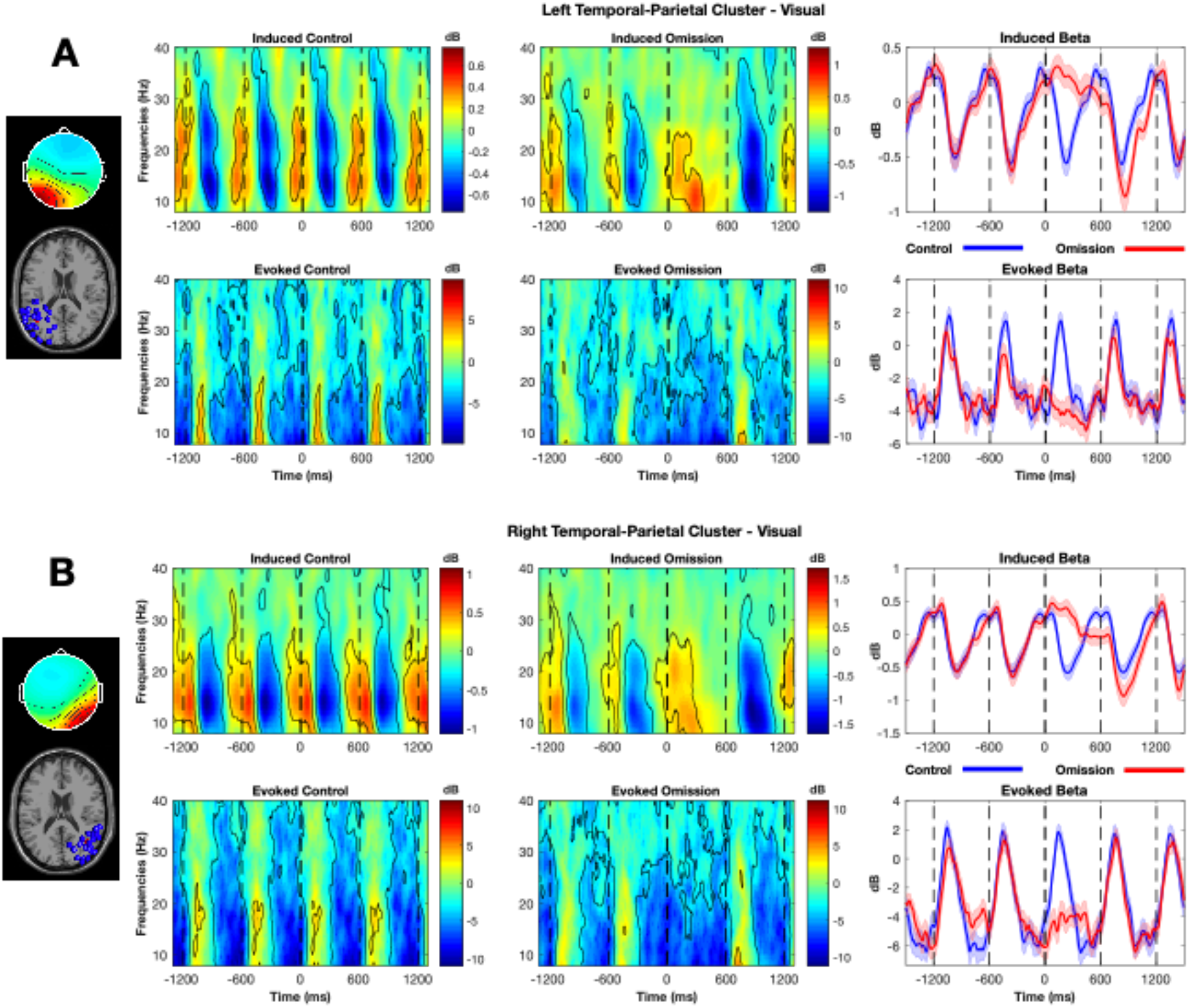
**I**nduced and evoked ERSP and beta band time frequency representations for the left (A) and right (B) temporal/parietal clusters for visual control and omission conditions. Significant time frequency values in the ERSP plot are outlined in black. Both clusters show induced beta power rising prior to the expected flash, and falling sharply after flash onset, and less sharply after the omission onset. Evoked beta power increases only in response to the flash, but not to the expected but omitted flash. Shaded bars in beta plots indicate standard error.

**Figure 5.**
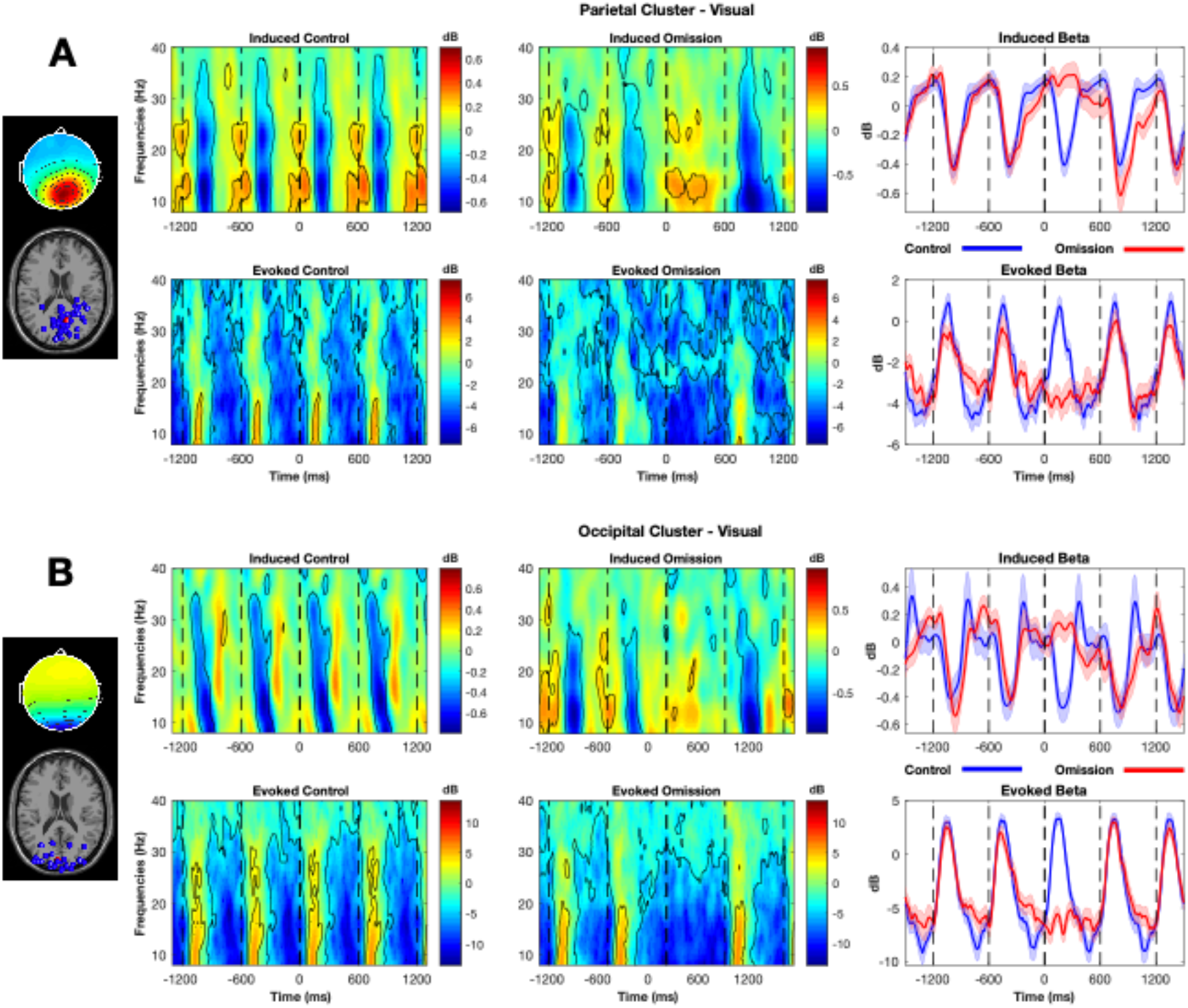
Induced and evoked ERSP and beta band time frequency representations for parietal (A) and occipital (B) clusters for visual control and omission conditions. Significant time frequency values in the ERSP plot are outlined in black. Shaded bars in beta plots indicate standard error. Parietal region induced beta power increases prior to expected stimulus onset, while occipital region induced beta power drops and rebounds sharply after stimulus onset. Both clusters show evoked beta power only increasing after stimulus onset.

**Figure 6.**
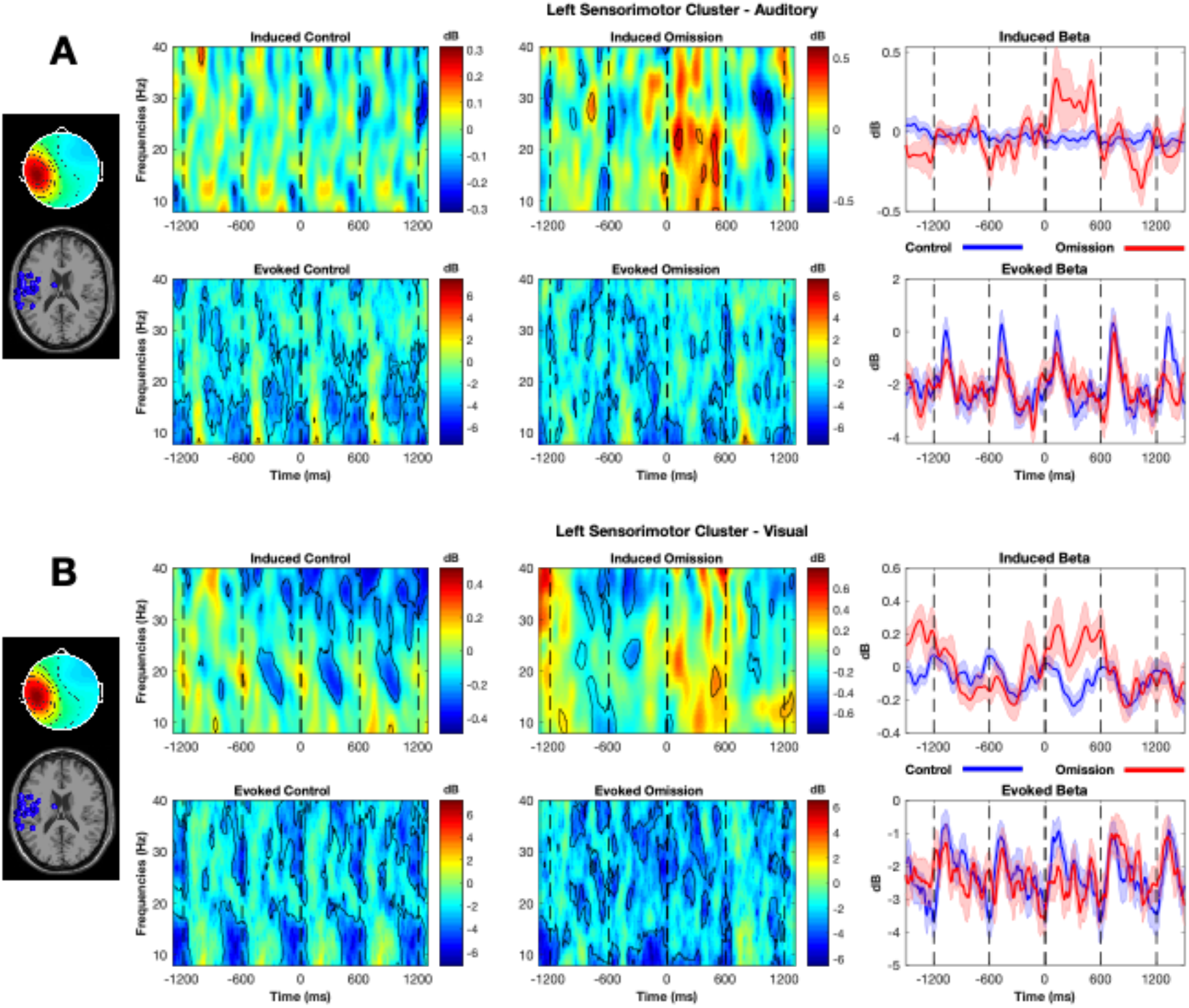
Induced and evoked ERSP and beta band time frequency representations for the left sensorimotor cluster for both auditory (A) and visual (B) conditions. Evoked beta power increases after both auditory tone onset and the expected but omitted tone onset. Visual evoked beta power is modulated by the flash onset, but not by the expected but omitted flash onset.

### Beta Band Slope Analysis

While significance testing in ERSP power can indicate significant power modulations in response to stimuli, we are interested in the dynamics of beta band activity following findings that indicate beta power rises to peak at the expected onset of an auditory tone, where the rate of the rise is dependent on the tempo of the stimuli (Fujioka et al., 2012). Since we hypothesized that rise in beta activity is related to the timing of the rhythmic stimuli, we would see beta power rise prior to the expected onset of the omitted stimuli. To test this hypothesis, 2 slopes were fitted in the averaged beta activity for each subject for each condition based on a least squares measure. The first slope started at −300 ms prior to stimulus or omission onset and ended at stimulus or omission onset (0 ms). Using −300 ms as the starting point was chosen as the halfway point between stimuli. Because there is considerable variation across subjects in slope activity, a second slope was fitted starting at the lowest measured activity between −300 and −100 ms and ending at stimulus or omission onset. To provide a third condition for comparison, we shuffled the ERSP data used to find slopes in the control condition for each subject at each channel, and for each component for each cluster, and then extracted beta band power and fitted slopes. Fitting a slope to the beta band extracted from the shuffled ERSP power results in an effective slope of 0, which we use to to compare the other slopes to. For the shuffled condition, ERSP power values along the entire time axis of each epoch of each frequency step were randomly shuffled 1000 times using the randperm function in Matlab. Beta band power for each time point was then extracted from resulting shuffled ERSPs the same as done with the non-shuffled ERSPs. Slopes were then fitted in the same way as with the non-shuffled data, except that instead of finding the minimum beta power between −300 and −100 ms for the shuffled condition, we used the same the same starting point used in the non-shuffled control condition for the corresponding subject or component.

Four sets of t-tests were used to determine if the fitted slope of beta activity prior to the onset of a tone or flash was equivalent to the fitted slope of beta activity prior to the expected but omitted onset of a tone or flash for both induced and evoked activity and for both the slopes the fitted from −300 ms to onset and for the slopes fitted to the trough between −300 and −100 ms and onset. FDR correction was used to correct for multiple comparisons for all t-tests using the method described in Benjamini and Hochberg (1995) with alpha set to 0.05. The first three analyses were performed using paired t-tests comparing: the slopes of the omission conditions to the slopes of the control conditions, the slopes of the control conditions to the slopes of the shuffled conditions, and the slopes of the omission conditions to the slopes of the shuffled conditions. If beta power is being modulated such that it shows anticipation of the stimulus rather than only reaction to the stimulus we would expect both the omission and control fitted slopes to be significantly different from the shuffled fitted slopes, and we would also expect the omission and non-omission fitted slopes to not be significantly different.

Showing that a fitted slope in the omission condition is not significantly different from the slope in the non-omission condition, yet significantly different from a flat slope is not sufficient to claim that the slopes in the omission and non-omission conditions are equivalent. This is because a comparison between significant results and nonsignificant results is not necessarily significant (Gelman & Stern, 2012). To assess the viability of the comparison between the two results, we applied a post-hoc comparison test as used in Abbott & Shahin (2018). The test calculated if the slope of the non-omission condition + the slope of the shuffled condition – 2 x the slope of the omission condition was significantly different from zero using a t-test with the same FDR correction as used for the other t-tests at each channel and each cluster.

The results of these tests at the electrode level show that only channel P8 meets the criteria for the 4 tests: p > 0.05 for the omission to non-omission slopes comparison, p < 0.05 for the comparisons of the non-omission to shuffled and omission to shuffled slopes, and p < 0.05 for the post hoc comparison test as applied to the slopes fitted to the between the trough of beta power and onset for induced beta. Additional channels met the first 3 criteria, but did not reach significance in the post-hoc test for the induced trough fitted slope for both visual and auditory conditions (Figure 8). No channels met these criteria for the slopes fitted at the fixed values between −300 ms and onset for the visual condition for induced or evoked beta. No auditory channels met the 4 criteria for any of the conditions.

At the cluster level in the visual modality 2 clusters plus the parent cluster met the criteria in induced activity for the slopes fitted to −300 to 0 ms: Clusters 9 (right temporal/parietal) and 8 (parietal). Cluster 6 (left temporal/parietal) met the criteria for three of the slope tests but not for the contrast (figures 4 & 5). No auditory clusters met the criteria for induced activity with a fixed slope. Slopes fitted to the trough (between −300 and −100 ms) and 0 ms for induced beta activity in the visual condition resulted in 5 clusters plus the parent cluster meeting the criteria for the 4 slope tests: Clusters 3 (midline central area), 5 (right frontal), 6 (left temporal/parietal), 8 (parietal region), and 9 (right temporal/parietal). Only cluster 7 (occipital) did meet the first 3 slope criteria in the visual modality for the trough fitted slope in induced activity. Cluster 8 (parietal) and the parent cluster met all 4 criteria for the auditory condition for trough fitted slopes to induced beta. All other clusters except 3 (midline central area) met the first 3 slope criteria for auditory induced beta trough fitted slopes.

Slopes fitted to evoked beta at the cluster level resulted in the parent cluster for both auditory and visual modalities, and cluster 2 (left sensorimotor) for the auditory modality meeting all 4 slope criteria for the trough fitted slopes (figure 6). Clusters 3 (midline central), 5 (right sensorimotor), and 8 (parietal) met the first 3 criteria for the trough fitted slope tests in both modalities. Clusters 6 (left temporal/parietal) and 1 (left frontal) in the auditory and visual modalities respectively met the first 3 slope criteria for the trough fitted slopes. No cluster met any of the necessary criteria in the slopes fitted between −300 and 0 ms to evoked beta activity. All slope measures and tests for the visual and auditory slopes can be found in the supplemental tables 1 (visual) and 2 (auditory)

### Evoked and Induced comparison

To further understand the different roles evoked and induced beta play in the temporal aspects of auditory and visual rhythm processing, we measured peak power and peak time in response to both present and omitted tones and flashes. To make the comparison ERSP power P was converted from dB to uV^2^ and normalized using the formula: P_norm_ = (P - P_min_) / (P_max_ - P_min_). This normalization conversion resulted in values between 0 and 1 and was applied to ERSP values for each individual component for each cluster after which beta power was extracted in the same manner as done for the slope analyses. Peak power and peak time were determined by finding the time and normalized power of the peak power between +/− 200 ms of the expected event onset. Paired t-tests were then run on each cluster as well as the parent cluster in order to determine the roles evoked and induced activity within each cluster. All t-tests used FDR correction to account for multiple comparisons. Test values presented here are for the parent cluster containing all components unless otherwise indicated. For a full listing of all test values and statistics for each cluster, refer to supplemental tables 3 (visual peak times), 4 (visual peak power), 5 (auditory peak times), and 6 (auditory peak power).

In the visual modality, evoked peak times for the control condition were generally after flash onset (M = 68.49 ms, SD = 122.18) and later than omission peak times (M = 11.04 ms, SD = 133.51); t(288) = 5.43, p = < 0.001. Visual induced peak times for the control condition tended to fall prior to onset (M = −12.95 ms, SD = 120.03), while omission peak times fell after expected onset (M = 28.74 ms, SD = 129.27); t(288) = −4.47, p = < 0.001. Both tests were also significant for clusters 3 (midline central) and 8 (parietal), with cluster 6 (left temporal/parietal) significant in induced activity and cluster 9 (right temporal/parietal) significant for evoked. The evoked control peak was significantly later than the induced control peak; t(288) = 8.06, p = < 0.001. This difference was also reflected in clusters 3 (midline central), 5 (right frontal), 6 (left temporal / parietal), 7 (occipital), 8 (parietal), and 9 (right temporal/parietal). Evoked and induced omission peak times were not significantly different in the parent cluster (t(288) = −1.67, p = 0.164), or any other cluster. To determine if the differences in control and omission peak times across induced and evoked activity were relative for each kind of activity, a further test compared the difference in evoked control and omission peak times (M = 57.44 ms, SD = 178) to the difference in induced control and omission peak times (M = −41.68 ms, SD = 158.43), revealing the relative shifts were significantly different; t(288) = 7.03, p = < 0.001. A significant relative difference was also seen in clusters 3 (midline central), 6 (left temporal/parietal), 7 (occipital), and 8 (parietal).

The same tests were run in the auditory on peak times, revealing that evoked auditory peak times for control (M = 10.64 ms, SD = 122.49) and omission (M = −2.23 ms, SD = 131.29) and induced auditory peak times for control (M = 0.35 ms, SD = 129.6) and omission (M = 6.7 ms, SD = 135.51) conditions were generally close to onset time and not significantly different from each other across all clusters and all tests except for cluster 2 (left sensorimotor), were evoked control peak time (M = 64.96, SD = 124.59) was significantly later than induced control peak time (M = −23.44, SD = 122.23); t(34) = 3.27, p = 0.006. The difference between evoked control and omission peak times (M = 35.38, SD = 167.02) and the difference between induced control and omission peak times (M = −45.42, SD = 151.1) was also found to be significant in cluster 2; t(34) = 2.39, p = 0.047.

Visual modality evoked control peak values (M = 0.631, SD = 0.141) were greater than evoked omission peak values (M = 0.366, SD = 0.163); t(288) = 20.04, p = <0.001. Similarly, visual modality induced control peak values (M = 0.753, SD = 0.117) were greater than induced omission peak values (M = 0.664, SD = 0.136), although to a lesser degree; t(288) = 9.58, p = <0.001. The comparison tests across visual omission and non-omission peak values within evoked and induced activity were significant for all clusters. Comparisons across evoked and induced peak values for visual beta indicated induced non-omission peaks were generally larger than evoked non-omission peaks; t(288) = −13, p = <0.001. This comparison was found to be significant for all clusters except cluster 7 (occipital). Comparisons across visual beta evoked and induced omission fitted peak values indicate induced omission peak values are greater than evoked omission peak values for the parent cluster; t(288) = −23.99, p = <0.001, and all other clusters. A comparison between the difference in evoked non-omission and omission peak power (M = 0.264, SD = 0.224) and the difference between induced non-omission and omission peak power (M = 0.089, SD = 0.157) indicated a greater relative difference was seen in evoked activity for the parent cluster (t(288) = 11.24, p = <0.001), as well as for clusters 3 (mid central), 5 (right frontal), 8 (parietal), and 9 (right temporal/parietal).

Running the same tests on auditory peak values show auditory evoked non-omission peak power (M = 0.592, SD = 0.125) was significantly greater than auditory evoked omission peak power (M = 0.442, SD = 0.146) for the parent cluster (t(288) = 13.24, p = <0.001), and all other clusters except for cluster 3 (midline central). Auditory induced non-omission peak power (M = 0.73, SD = 0.099) was slightly larger than auditory induced omission peak power (M = 0.677, SD = 0.124), and significantly so for the parent cluster (t(288) = 5.72, p = <0.001), as well as for clusters 3 (midline central), 7 (occipital), and 9 (right temporal/parietal). A comparison across auditory evoked and induced non-omission peak power reveals induced non-omission peak power is significantly greater in the parent cluster (t(288) = −15.65, p = <0.001), as well as in all other clusters except cluster 6 (left frontal). Auditory induced omission peak power was found significantly larger in the parent cluster (t(288) = −20.27, p = <0.001), as well as all other clusters. Comparing the difference in evoked non-omission and omission peak power (M = 0.15, SD = 0.193) and the difference between induced non-omission and omission peak power (M = 0.053, SD = 0.156) revealed a greater relative difference in evoked activity that was significant in parent cluster (t(288) = 6.51, p = <0.001), as well as for clusters 2 (left sensorimotor), 4 (right sensorimotor), 5 (right frontal), and 8 (parietal).

## Discussion

### Summary of Results

Using a cluster based approach to describe network-level beta band activity, we described predictive timing in a modality-specific way. Analyses on the slopes of beta activity from the parent clusters reveal evidence for both induced and evoked predictive timing in auditory and visual modalities at the global level. A look at the slopes of beta activity from individual clusters indicates evidence of induced predictive timing in the visual modality in posterior regions: left and right temporal/parietal clusters, and parietal cluster; the midline central cluster, and from the right frontal cluster. Slope based evidence for induced predictive timing in the auditory modality was found in the parietal cluster. Cluster specific evidence of evoked predictive timing in slope measures was seen only in the auditory modality, and only in the left sensorimotor cluster.

Based on previous results from Snyder & Large (2005) we expected evoked beta peak power to be significantly lower for omission events compared to tone or flash events, and we expected there to be no significant difference in induced beta peak power between omission events and tone or flash events. This pattern was seen much more prominently in the auditory modality, specifically in the parietal, left and right sensorimotor, left and right frontal, and left temporal/parietal clusters. A significant difference would additionally be expected between how much evoked beta peak power shifted between non-omission and omission conditions and how much induced beta power shifted between non-omissions and omissions. This significant difference was replicated in several clusters: the parietal cluster, left and right sensorimotor clusters, and the right frontal cluster, thus providing strong evidence for auditory induced beta playing a predictive role in networks of those regions. There were a few differences in the peak times in auditory beta across both induced and evoked activity and conditions. The significant shift in peak time from tone to omitted tone between induced and evoked beta for the right sensorimotor cluster follows the expected pattern of induced beta peaking later in response to an omitted tone than in response to a non-omitted tone. The evoked beta peaked earlier in response to an omitted tone than in response to a non-omitted tone. While not significant, we find it interesting that the opposite pattern with beta peak time appears in the left sensorimotor cluster: induced beta peaked slightly earlier in response to omitted tones than in response to tones, yet evoked beta peaked slightly later in response to the omitted tones than in response to the tones. This is in concordance with what would be expected if evoked beta was playing a predictive role, and when taken in conjunction with the slope evidence of predictive evoked activity in the left sensorimotor cluster suggests the existence of significant hemispheric differences in auditory rhythm processing mechanisms.

Differences in evoked and induced beta power in response to visual non-omissions and omissions did not provide clear evidence of predictive beta as seen in the auditory case, except for in the shift of peak power between evoked and induced activity from flash to flash omission in the parent, parietal, midline central, right frontal, and right temporal/parietal clusters. Interestingly, a look at differences in peak times does provide stronger evidence suggesting separate roles for evoked and induced beta for the parietal, right and left temporal/parietal, and occipital clusters. In these clusters the evoked beta peak came earlier in response to omitted flashes than to non-omitted flashes, while induced beta peaked later in response to omitted flashes than to non-omitted flashes, which is what would be expected if induced beta activity was playing a predictive role, while evoked beta was only responsive to stimuli. Taken together with the slope results, we interpret these findings as evidence of induced beta playing a predictive role in visual rhythm perception similar to that reported in previous studies for auditory induced beta (Fujioka et al., 2009, 2012, 2015; Snyder & Large 2005).

### Predictive Beta band activity

Beta modulation has been shown to play a role in a wide range of activities including top down control on sensorimotor systems (Engel & Fries, 2010; Arnal et al., 2011; Picazio et al, 2014; Haegens & Golumbic, 2018), facilitating long-range communication between cortical regions (Kopell et al., 2000; Kilavik et al., 2013) such as between sensorimotor and peripheral areas (Fujioka et al., 2015), and is suggested to play a role in encoding temporal intervals (Wiener et al., 2016). Beta band activity also correlates with motor behavior, with power attenuation just before and during movements (See Kilavik et al., 2013 for review). Considering the suggested role the motor cortex has in timing and predictive processing (Schubotz et al., 2000; Patel & Iversen, 2014), the role of beta in imposing general top down control, and its role in facilitating communication with sensorimotor peripheral systems, it is not surprising that beta activity appears to play a role in rhythm perception and prediction.

Beyond the link to sensorimotor behavior, beta activity is known to play a role in auditory rhythm perception. Frontocentral induced beta and gamma modulation occurs with the onset of rhythmic events and can be seen at the expected onset of an omitted event (Snyder & Large, 2005). Fujioka et al. (2012) found that beta power arising from the auditory cortices increases before tone onset in an isochronous rhythm at a rate dependent on the tempo of the rhythm, and attenuates following the tone at a constant rate not dependant on the tempo of the rhythm. Beta activity has also been seen to play a role in maintaining beat and meter structure (Fujioka et al., 2015). Consistent with these findings, we find evidence of auditory induced beta power peaking in anticipation of both tones and omitted tones, with the strongest evidence coming from the parietal, left and right sensorimotor, and right frontal clusters. Because the source of neural activations are more difficult to localize using EEG than MEG, some caution is needed in interpreting the location of these sources. However, given other findings suggesting predictive induced beta arising from fronto-central regions using EEG (Snyder & Large 2005), and from the auditory cortices, sensorimotor cortices, and parietal cortices using MEG (Fujioka et al., 2012, 2015), we believe the regions indicated by the cluster locations are reasonable interpretations of the source of the predictive beta we measured. It is of note that we did not find evidence of predictive beta that we could tie clearly to the auditory cortex. This may be a limitation of the cluster approach we used with the independent components, but it has also been put forth that signals arising from the auditory cortex are more suited to being measured by MEG than EEG (Destoky et al., 2019).

When looking at beta modulation in the visual domain, we see a beta power increase at the expected onset of an omitted flash in multiple clusters. Comparing beta modulation in anticipation of the visual onset between the omission and non-omission conditions shows induced beta power increasing prior to onset, followed by a sharp power drop-off, but only after flash onset, and not following omission onset. While we expected to find predictive beta activity in the visual domain, it was surprising to see evidence of predictive induced beta modulated more clearly and across more clusters in the visual domain than in the auditory domain because the timing aspects of rhythm perception in the auditory domain are thought to be more precise as evinced by less variability in auditory SMS compared to visual SMS (Repp 2005, Repp & Su 2013). We suggest this discrepancy between auditory and visual beta modulation is due a combination of factors. The most important factor being the size differential between the visual and auditory cortices; the visual cortex is much larger than the auditory cortex, and so processing of visual stimuli involves more cortical neurons resulting in more neural activity measured at the scalp than auditory cortex would produce. Compounding this is the suggestion previously mentioned that auditory signals are more suited to measurement from MEG than from EEG (Destoky et al., 2019), resulting in a comparatively reduced measurement of beta modulated by auditory rhythms.

The clusters that show evidence of predictive beta activity for the visual modality do not perfectly overlap with what is seen in the auditory modality. In the sensorimotor clusters, we only find evidence of auditory predictive beta in bilateral sensorimotor clusters, and not visual predictive beta. There is evidence of visual predictive beta in the midline cluster, which contains dipoles localized to the premotor regions. This may indicate motor system involvement and would be inline with research suggesting the medial premotor region plays a role in predictive timing in primates across sensory modalities (Merchant et al., 2013). However, this begs the question of why the same activity was not seen in the auditory modality if premotor timing activity is not modality specific. A possible explanation is given by work reporting that a greater number of cells in the primate SMA respond to visual timing cues than to auditory timing cues (Merchant et al., 2015), although it is not clear if this finding extends to humans or if it is specific to the primates involved in that study. It is also of interest that we find predictive visual induced beta activity from the slope analysis in left and right temporal/parietal junction and parietal clusters, but not in the occipital cluster. Given the difficulty in localizing sources with EEG, and the component distribution of the four posterior clusters, it is likely the left and right temporal/parietal and parietal clusters contain activity arising from cortical patches within the occipital cortex. Considering the distribution of components, and the faster rebound in induced beta power in the occipital cluster (figure 5b), we consider it likely that activity from early processing areas of the visual cortex (e.g. V1) are more strongly represented in the occipital cluster than the surrounding posterior clusters. This however cannot be confirmed with the spatial limitations of EEG, and will require a methodology with greater spatial precision to test.

While beta power modulation in response to visual rhythmic flashes has been seen before (Saleh et al., 2010, Meijer et al., 2016), to our knowledge this is the first time it has been shown predicting the onset of an omitted event. However, it has been questioned whether beta modulation is even related to temporal prediction at all (Meijer et al., 2016). Meijer et al., (2016) investigated beta activity with a rhythmic visual task and found beta power modulation in response to isochronous visual rhythms of different tempi (IOI’s of 1050, 1350, 1650 ms), yet the rate of beta power modulation was the same regardless of the tempo used. This is different from what was found by Fujioka et al., (2012) in their study of auditory beta modulation, where the rate of beta power prior to tone onset was modulated by the tempo of the rhythm. Meijer et al., (2016) interpreted their result as evidence that beta activity is not playing an entraining role in the visual system, suggesting instead that the beta peaks seen may be caused by rebounding activity in response to the flash, peaking roughly 900 ms after event onsets. The current study provides the contrary evidence, and suggests that beta modulation may be playing a role in prediction of the onset of visual events, since the beta modulation during the omission could not be in response to any event, and instead must be responding to the timing of the expected onset of the flash. Induced beta peaks less than 50 ms after the omission onset, or 650 ms after the onset of the prior stimulus (figure 4), which is much earlier than would be expected for beta power rebound in response to the flash event, as described by Meijer et al. (2016). We suggest the reason for the discrepancy between Meijer et al.’s (2016) findings and those findings reported here may be due to their use of relatively slow tempi compared to the 600 ms IOI of this study. There is evidence that sub-second timing and supra-second timing use different networks (see Wiener et al, 2010 for a review). We therefore suggest beta synchronization may only be playing a predictive role in the sub second time scale., the task used in the Meijer et al. (2016) study was much more complicated than simply attending to the timing of the rhythms as in our task, and demanded more attention and possibly competing resources.

### Contribution of the motor system

Previous studies have described induced beta modulation to auditory rhythms arising from sensorimotor cortices (Fujioka et al., 2012, 2015). There is also evidence that auditory timing appears to rely on motor cortex (Janata et al., 2012; Repp and Su, 2013; Iversen and Balasubramaniam, 2016; Ross et al., 2016a, 2016b) and motor networks with nodes in the parietal lobes, cerebellum, and basal ganglia (Repp & Su, 2013; Patel & Iversen, 2014; Levitin et al., 2018). This motor network activity could indicate that the motor system is playing an important role in predicting the timing of events in auditory rhythms, often discussed in the context of evolution of social activities such as dance and language. (Fitch, 2016; Iversen, 2016; Patel, 2006). The auditory beta modulation from the sensorimotor clusters we present here is consistent with the narratives of the previous literature on the involvement of the motor system for auditory timing. This can be contrasted with our findings from the visual system where there is no evidence of predictive beta timing in the bilateral sensorimotor clusters, and instead evidence in the mid-central cluster that may be related activity arising from the SMA.

In the auditory modality, we found evoked predictive beta timing activity in the left sensorimotor cluster (figure 6a), yet we found evidence of induced predictive timing activity in the right sensorimotor cluster (figure 7a). The asymmetrical beta activity seen in the two sensorimotor clusters specific to the auditory conditions suggests hemispheric specialization specific to auditory processing. A recent meta-analysis on neural activation during music listening shows consistent MRI activation in the right but not left primary motor cortex during music listening tasks (Gordon et al., 2018). Interestingly, they found that studies that asked the subjects to move a body part while listening elicited stronger activity in the right primary motor cortex than studies using passive listening tasks. Others describe a left hemisphere role (Pollok, Rothkegel, Schnitzler, Paulus, & Lang, 2008) or non-motor-dominant hemisphere role (Kaulmann, Hermsdörfer, & Johannsen, 2017; Yadav & Sainburg, 2014) for motor timing. Similarly, for language perception there appears to be hemispheric specialization in the auditory cortices, with the left hemisphere specialized in temporal changes and the right hemisphere in spectral changes (Zatorre et al., 1992; Zatorre & Belin, 2001). Specifically, it has been shown that activity in the left anterolateral superior temporal sulcus (STS) corresponds to processing of temporal aspects of speech perception, while perception of spectral features of speech are associated with the same structure in the right hemisphere (Obleser et al., 2008). Our results support bilateral motor contributions to auditory timing, although the mechanism that results in predictive evoked activity in the left hemisphere and predictive induced beta activity in the right hemisphere may be distinct.

**Figure 7.**
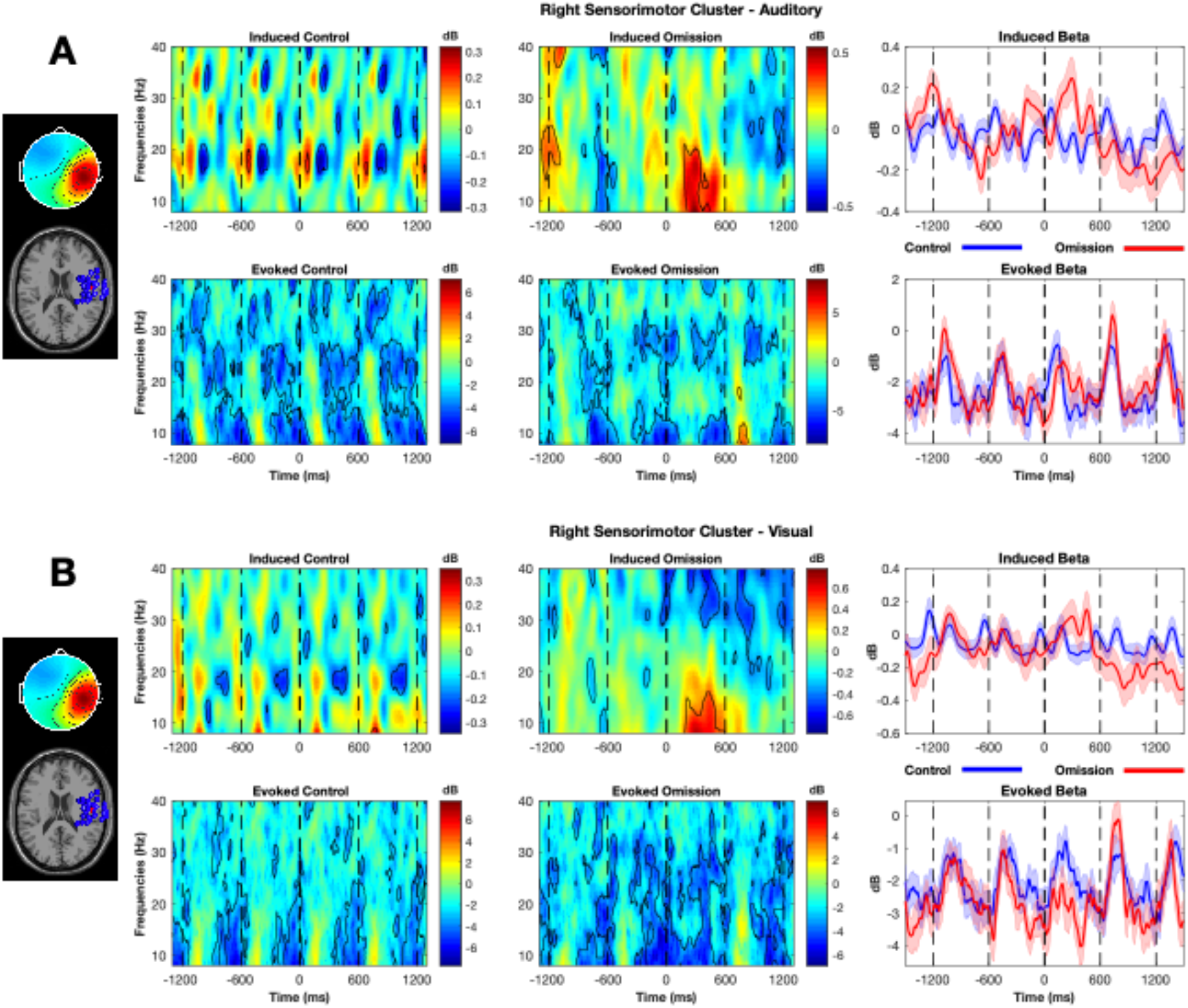
Induced and evoked ERSP and beta band time frequency representations for the right sensorimotor cluster for both auditory (A) and visual (B) conditions. Both auditory and visual conditions appear to show stimulus modulated induced beta power that is not modulated by the expected stimulus onset. Evoked beta for both auditory and visual conditions increases after stimulus onset, and appears to also increase after the omission onset, but not significantly.

**Figure 8.**
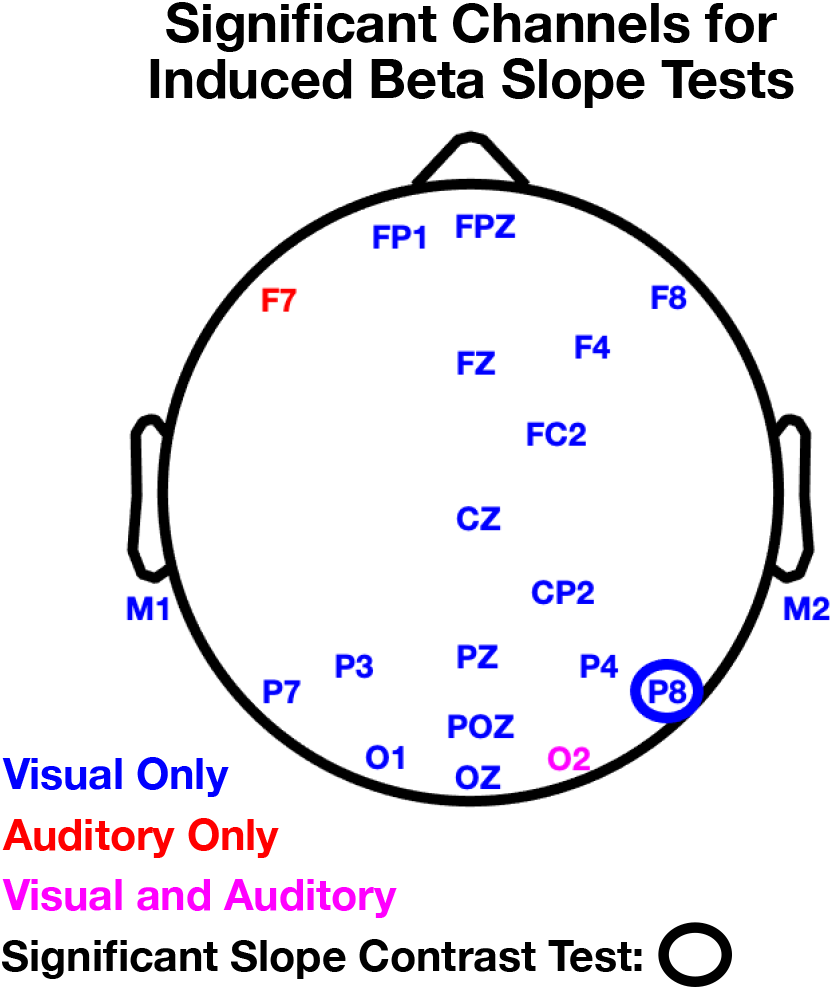
Significant channels for the induced beta tests to slopes fitted from the trough of beta power between −300 and −100 ms to the event onset at 0 ms. Channels labeled had p > 0.05 for the omission to control slopes comparison, and p < 0.05 for the comparisons of the control to shuffled and omission to shuffled slopes. The circled channel (P8) indicates p < 0.05 for the post hoc comparison test as applied to the slopes fitted to the between the trough of beta power and onset for induced beta.

### Limitations & Future Directions

The current study reveals that timing and prediction for visual rhythm perception could employ non-motor networks. We cannot say what role, if any, the motor system plays in visual timing. A closer look at the connections between visual and motor systems is needed to elucidate the issue. Using flashing visual rhythms as opposed to moving visual rhythms may elicit a different picture of activation as the visual system is better tuned to discerning temporal information when movement is present (Hove et al, 2013b).

Another limitation of the current study is that we did not use multiple tempi. Having only one tempo makes it unclear how much the change in time course of neural activations is related to the tempo. Using multiple rhythms with different tempi would allow for a clearer differentiation between tempo dependent aspects of timing. If those tempi spanned both sub-second and supra-second timing it would also provide insight to the temporal limits to the mechanisms in visual rhythm perception.

Although we see frequency band specific oscillatory modulation during rhythm perception, caution should be used in assuming this is the brain’s mechanism of timing. There is evidence for multiple mechanisms for timing (for review see: Wiener et al., 2010; Wiener & Kanai, 2016; Comstock, Hove, & Balasubramaniam, 2018), and here we describe one reflection of these processes. Oscillatory dynamics likely reflect more broadly the mechanism for spreading information between or across networks, and timing perception is only a subset of neural communication happening during these tasks.

Additional investigation is needed into the differences seen between left and right motor contributions to auditory timing. While the differences suggest possible functional lateralization in auditory rhythm perception, it is unclear if those differences are driven by handedness (Kaulmann, Hermsdörfer, & Johannsen, 2017; Yadav & Sainburg, 2014) or other factors (Pollok, Rothkegel, Schnitzler, Paulus, & Lang, 2008). Future studies are needed to look more closely at specific hemispheric contributions.

Finally, the inherent low spatial resolution of EEG limits how confidently we can draw conclusions about neural sources. We describe broad cortical source regions/networks in lieu of more focal sources with respect to this methodological limitation, but argue that the ICA-based cluster analysis leads to reasonable spatial and functional grouping of neural activity likely from common sources. That being said, we cannot speak with certainty about the exact cortical sources of the activity we describe. A method with better spatial resolution that retains fine temporal resolution, such as MEG or ECoG, would provide better source resolution for predictive rhythm perception networks.

### Conclusion

We investigated the mechanisms of prediction for auditory and visual rhythms using an omission paradigm. Results show induced beta activity predicting the expected onset of visual rhythmic events bilaterally in temporal/parietal clusters, in a dorsal medial cluster, a parietal cluster, and a right hemisphere frontal cluster. We also show induced beta activity predicting the expected onset of rhythmic auditory events bilaterally in sensorimotor clusters, in a parietal cluster, and in a right hemisphere frontal cluster. We additionally present evidence for evoked auditory predictive timing in a left motor cluster. Our results support theories of predictive timing in both visual and auditory modalities, that can be observed in beta band oscillatory activity. Our results also support, using a cluster based approach, that visual and auditory prediction for rhythmic events may be subserved by modality-specific cortical networks, although they do not rule out the possibility that both auditory and visual networks are subserved by a common subcortical network. These findings also suggest that auditory timing may involve hemisphere specific activity, and reliance on motor networks.

## Autor Contributions

DCC and RB authors conceived and designed this study together. DCC ran all participants and conducted the original analyses. DCC and JMR conducted follow-up analysis. DCC, JMR, and RB co-wrote the paper.

## Acknowledgements

The work was supported by NSF grant BCS 1626505.

## Accessibility

EEG data, code and stimuli from this study are available at https://openneuro.org/datasets/ds002218. The EEGLAB (v14.1.1) tools that were used to analyze all the data can be downloaded from the Swartz Center for Computational Neuroscience website: https://sccn.ucsd.edu/eeglab/download.php. Paradigm software (ver. 2.5.0.68) was used to present the instructions, stimuli, and to sync the start of each trial with the EEG data.

## Abbreviations

ASR: artifact source reconstruction
cm: centimeters
Db: decibel
EEG: electroencephalography
ECoG: Electrocortocography
ERSP: event related spectral perturbation
Hz: hertz
ICA: independent component analysis
IOI: interonset interval
ITC: inter-trial coherence
MAD: mean absolute deviation
MEG: magnetoencephalography
MRI: magnetic resonance imaging
ms: milliseconds
SD: standard deviation
SMA: supplementary motor area
SMS: sensorimotor synchronization
STS: superior temporal sulcus

## Supplemental Materials

**Supplemental Table 1:**
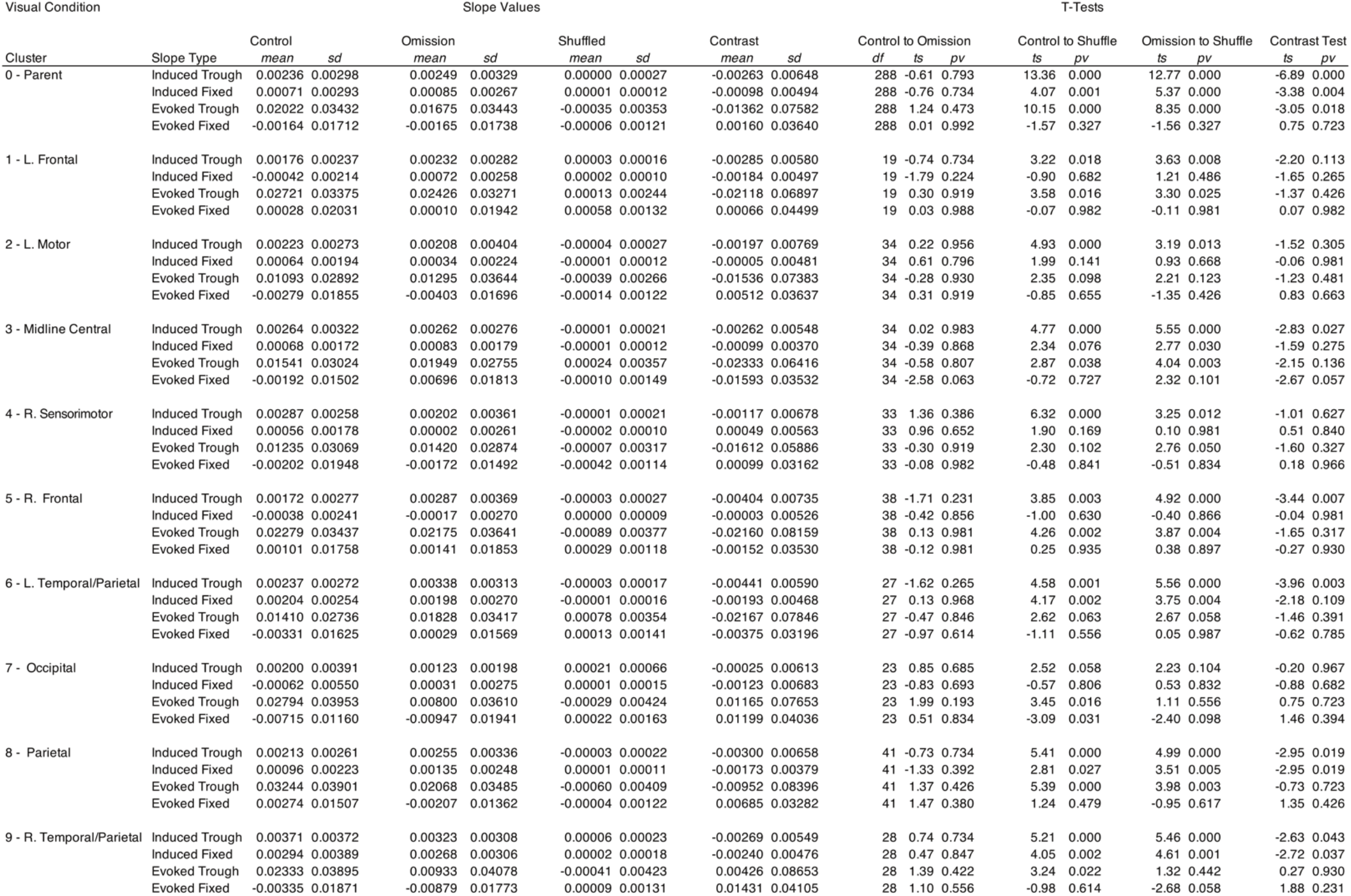
Visual Slope statistics for each cluster for induced and evoked slopes. Contrast values are calculated as the control slope + the shuffle slope – (2 * omission slope). The contrast test compares the contrast values to zero. Degrees of freedom were the same for each of four t-tests on the same row. P-values displayed are FDR corrected.

**Supplemental Table 2.** Auditory Slope statistics for each cluster for induced and evoked slopes. Contrast values are calculated as the control slope + the shuffle slope – (2 * omission slope). The contrast test compares the contrast values to zero. Degrees of freedom were the same for each of four t-tests on the same row. P-values displayed are FDR corrected.

| Auditory Condition |  | Slope Values |  |  |  |  |  |  |  | T-Tests |  |  |  |  |  |  |  |  |
| --- | --- | --- | --- | --- | --- | --- | --- | --- | --- | --- | --- | --- | --- | --- | --- | --- | --- | --- |
| Cluster | Slope Type | Control |  | Omission |  | Shuffled |  | Contrast |  | Control to Omission |  |  | Control to Shuffle |  | Omission to Shuffle |  | Contrast Test |  |
|  |  | mean | sd | mean | sd | mean | sd | mean | sd | df | ts | pv | ts | pv | ts | pv | ts | pv |
| 0 - Parent | Induced Trough | 0.00186 | 0.00308 | 0.00202 | 0.00335 | 0.00001 | 0.00021 | -0.00216 | 0.00701 | 288 | -0.62 | 0.793 | 10.20 | 0.000 | 10.13 | 0.000 | -5.24 | 0.000 |
|  | Induced Fixed | 0.00008 | 0.00189 | 0.00009 | 0.00209 | 0.00001 | 0.00009 | -0.00009 | 0.00476 | 288 | -0.04 | 0.981 | 0.67 | 0.761 | 0.67 | 0.761 | -0.32 | 0.901 |
|  | Evoked Trough | 0.02290 | 0.03363 | 0.02033 | 0.03322 | -0.00004 | 0.00306 | -0.01780 | 0.07456 | 288 | 0.92 | 0.618 | 11.51 | 0.000 | 10.40 | 0.000 | -4.06 | 0.001 |
|  | Evoked Fixed | 0.00109 | 0.01779 | -0.00055 | 0.01848 | 0.00015 | 0.00117 | 0.00233 | 0.04133 | 288 | 1.07 | 0.556 | 0.90 | 0.628 | -0.64 | 0.769 | 0.96 | 0.614 |
| 1 - L. Frontal | Induced Trough | 0.00105 | 0.00424 | 0.00163 | 0.00232 | 0.00007 | 0.00040 | -0.00214 | 0.00555 | 19 | -0.58 | 0.806 | 0.96 | 0.655 | 2.93 | 0.029 | -1.73 | 0.240 |
|  | Induced Fixed | -0.00084 | 0.00277 | 0.00097 | 0.00238 | -0.00001 | 0.00012 | -0.00279 | 0.00590 | 19 | -2.05 | 0.141 | -1.35 | 0.397 | 1.86 | 0.197 | -2.12 | 0.129 |
|  | Evoked Trough | 0.00927 | 0.02620 | 0.02828 | 0.03242 | -0.00014 | 0.00185 | -0.04742 | 0.07493 | 19 | -1.86 | 0.250 | 1.66 | 0.324 | 3.92 | 0.008 | -2.83 | 0.054 |
|  | Evoked Fixed | 0.00341 | 0.01589 | 0.00328 | 0.01881 | -0.00030 | 0.00084 | -0.00345 | 0.04142 | 19 | 0.02 | 0.988 | 1.03 | 0.593 | 0.86 | 0.655 | -0.37 | 0.897 |
| 2 - L. Sensorimotor | Induced Trough | 0.00150 | 0.00270 | 0.00189 | 0.00444 | -0.00002 | 0.00019 | -0.00230 | 0.00862 | 34 | -0.50 | 0.841 | 3.30 | 0.011 | 2.53 | 0.050 | -1.58 | 0.277 |
|  | Induced Fixed | -0.00001 | 0.00169 | 0.00006 | 0.00223 | 0.00000 | 0.00007 | -0.00012 | 0.00510 | 34 | -0.12 | 0.969 | -0.04 | 0.981 | 0.14 | 0.968 | -0.13 | 0.968 |
|  | Evoked Trough | 0.01107 | 0.02256 | 0.03143 | 0.03536 | 0.00001 | 0.00242 | -0.05177 | 0.07661 | 34 | -2.74 | 0.050 | 2.92 | 0.035 | 5.25 | 0.000 | -4.00 | 0.004 |
|  | Evoked Fixed | 0.00308 | 0.01653 | 0.00411 | 0.01442 | -0.00015 | 0.00110 | -0.00529 | 0.03070 | 34 | -0.30 | 0.919 | 1.15 | 0.537 | 1.77 | 0.267 | -1.02 | 0.593 |
| 3 - Midline Central | Induced Trough | 0.00183 | 0.00354 | 0.00215 | 0.00391 | 0.00003 | 0.00020 | -0.00244 | 0.00714 | 34 | -0.46 | 0.847 | 3.02 | 0.019 | 3.17 | 0.014 | -2.02 | 0.137 |
|  | Induced Fixed | -0.00005 | 0.00140 | -0.00009 | 0.00231 | 0.00001 | 0.00009 | 0.00015 | 0.00481 | 34 | 0.09 | 0.981 | -0.28 | 0.929 | -0.27 | 0.929 | 0.18 | 0.968 |
|  | Evoked Trough | 0.02864 | 0.03863 | 0.01893 | 0.03400 | 0.00083 | 0.00297 | -0.00839 | 0.07216 | 34 | 1.24 | 0.481 | 4.25 | 0.002 | 3.09 | 0.025 | -0.69 | 0.747 |
|  | Evoked Fixed | -0.00319 | 0.01843 | 0.00039 | 0.01859 | 0.00027 | 0.00103 | -0.00369 | 0.03589 | 34 | -0.97 | 0.614 | -1.09 | 0.556 | 0.04 | 0.988 | -0.61 | 0.785 |
| 4 - R. Sensorimotor | Induced Trough | 0.00186 | 0.00293 | 0.00118 | 0.00256 | 0.00003 | 0.00022 | -0.00047 | 0.00592 | 33 | 1.02 | 0.625 | 3.72 | 0.004 | 2.62 | 0.042 | -0.46 | 0.847 |
|  | Induced Fixed | 0.00027 | 0.00203 | 0.00038 | 0.00156 | -0.00002 | 0.00010 | -0.00051 | 0.00403 | 33 | -0.23 | 0.952 | 0.83 | 0.689 | 1.48 | 0.323 | -0.74 | 0.734 |
|  | Evoked Trough | 0.01731 | 0.02658 | 0.00938 | 0.02251 | 0.00013 | 0.00285 | -0.00133 | 0.05211 | 33 | 1.31 | 0.442 | 3.67 | 0.008 | 2.43 | 0.087 | -0.15 | 0.977 |
|  | Evoked Fixed | 0.00160 | 0.01797 | -0.00053 | 0.01891 | 0.00002 | 0.00127 | 0.00268 | 0.04903 | 33 | 0.39 | 0.896 | 0.51 | 0.834 | -0.17 | 0.972 | 0.32 | 0.919 |
| 5 - R. Frontal | Induced Trough | 0.00286 | 0.00353 | 0.00231 | 0.00389 | -0.00011 | 0.00029 | -0.00187 | 0.00837 | 38 | 0.66 | 0.771 | 5.11 | 0.000 | 3.94 | 0.002 | -1.39 | 0.366 |
|  | Induced Fixed | -0.00017 | 0.00179 | -0.00039 | 0.00242 | 0.00000 | 0.00010 | 0.00060 | 0.00534 | 38 | 0.42 | 0.856 | -0.59 | 0.803 | -0.98 | 0.641 | 0.70 | 0.757 |
|  | Evoked Trough | 0.02024 | 0.03283 | 0.02181 | 0.02620 | 0.00065 | 0.00294 | -0.02273 | 0.06200 | 38 | -0.23 | 0.945 | 3.77 | 0.005 | 5.01 | 0.000 | -2.29 | 0.102 |
|  | Evoked Fixed | 0.00120 | 0.01681 | 0.00024 | 0.02091 | -0.00011 | 0.00136 | 0.00061 | 0.04832 | 38 | 0.20 | 0.961 | 0.49 | 0.840 | 0.11 | 0.981 | 0.08 | 0.982 |
| 6 - L. Temporal/Parietal | Induced Trough | 0.00191 | 0.00314 | 0.00238 | 0.00320 | 0.00004 | 0.00026 | -0.00281 | 0.00710 | 27 | -0.56 | 0.811 | 3.12 | 0.017 | 3.79 | 0.004 | -2.10 | 0.126 |
|  | Induced Fixed | 0.00022 | 0.00192 | 0.00023 | 0.00171 | -0.00001 | 0.00009 | -0.00025 | 0.00380 | 27 | -0.02 | 0.983 | 0.63 | 0.793 | 0.75 | 0.734 | -0.35 | 0.883 |
|  | Evoked Trough | 0.01678 | 0.02840 | 0.02216 | 0.04056 | -0.00078 | 0.00331 | -0.02832 | 0.09018 | 27 | -0.53 | 0.833 | 3.18 | 0.025 | 3.02 | 0.032 | -1.66 | 0.317 |
|  | Evoked Fixed | -0.00053 | 0.01547 | -0.00088 | 0.02210 | -0.00023 | 0.00125 | 0.00100 | 0.04488 | 27 | 0.07 | 0.982 | -0.10 | 0.981 | -0.16 | 0.974 | 0.12 | 0.981 |
| 7 - Occipital | Induced Trough | 0.00152 | 0.00239 | 0.00186 | 0.00296 | 0.00001 | 0.00023 | -0.00219 | 0.00627 | 23 | -0.46 | 0.847 | 3.18 | 0.017 | 2.95 | 0.026 | -1.71 | 0.240 |
|  | Induced Fixed | 0.00039 | 0.00205 | -0.00015 | 0.00192 | 0.00002 | 0.00009 | 0.00071 | 0.00400 | 23 | 1.03 | 0.625 | 0.87 | 0.682 | -0.44 | 0.851 | 0.87 | 0.682 |
|  | Evoked Trough | 0.03113 | 0.03413 | 0.01391 | 0.03010 | -0.00045 | 0.00305 | 0.00287 | 0.06284 | 23 | 2.03 | 0.182 | 4.47 | 0.002 | 2.41 | 0.098 | 0.22 | 0.949 |
|  | Evoked Fixed | 0.00661 | 0.01898 | -0.00177 | 0.01523 | 0.00032 | 0.00130 | 0.01047 | 0.03336 | 23 | 1.83 | 0.250 | 1.60 | 0.327 | -0.66 | 0.768 | 1.54 | 0.362 |
| 8 - Parietal | Induced Trough | 0.00135 | 0.00211 | 0.00210 | 0.00304 | 0.00003 | 0.00017 | -0.00283 | 0.00622 | 41 | -1.39 | 0.366 | 3.94 | 0.002 | 4.41 | 0.001 | -2.94 | 0.019 |
|  | Induced Fixed | 0.00012 | 0.00167 | -0.00003 | 0.00187 | -0.00001 | 0.00007 | 0.00017 | 0.00428 | 41 | 0.37 | 0.879 | 0.51 | 0.840 | -0.06 | 0.981 | 0.26 | 0.938 |
|  | Evoked Trough | 0.03394 | 0.04097 | 0.02020 | 0.03605 | -0.00032 | 0.00310 | -0.00678 | 0.08332 | 41 | 1.63 | 0.321 | 5.46 | 0.000 | 3.67 | 0.006 | -0.53 | 0.833 |
|  | Evoked Fixed | 0.00076 | 0.01546 | -0.00282 | 0.01891 | -0.00031 | 0.00095 | 0.00609 | 0.04178 | 41 | 0.92 | 0.621 | 0.45 | 0.858 | -0.86 | 0.655 | 0.94 | 0.617 |
| 9 - R. Temporal/Parietal | Induced Trough | 0.00258 | 0.00313 | 0.00270 | 0.00285 | 0.00002 | 0.00027 | -0.00280 | 0.00678 | 28 | -0.14 | 0.968 | 4.45 | 0.001 | 5.08 | 0.000 | -2.22 | 0.101 |
|  | Induced Fixed | 0.00057 | 0.00188 | 0.00021 | 0.00227 | 0.00000 | 0.00010 | 0.00016 | 0.00534 | 28 | 0.60 | 0.798 | 1.64 | 0.264 | 0.48 | 0.847 | 0.17 | 0.968 |
|  | Evoked Trough | 0.03172 | 0.03733 | 0.01840 | 0.03940 | -0.00034 | 0.00345 | -0.00542 | 0.08205 | 28 | 1.43 | 0.398 | 4.66 | 0.001 | 2.56 | 0.069 | -0.36 | 0.900 |
|  | Evoked Fixed | -0.00203 | 0.02424 | -0.00699 | 0.01736 | -0.00015 | 0.00132 | 0.01180 | 0.04222 | 28 | 0.90 | 0.633 | -0.42 | 0.881 | -2.12 | 0.148 | 1.51 | 0.371 |

**Supplemental Table 3.**
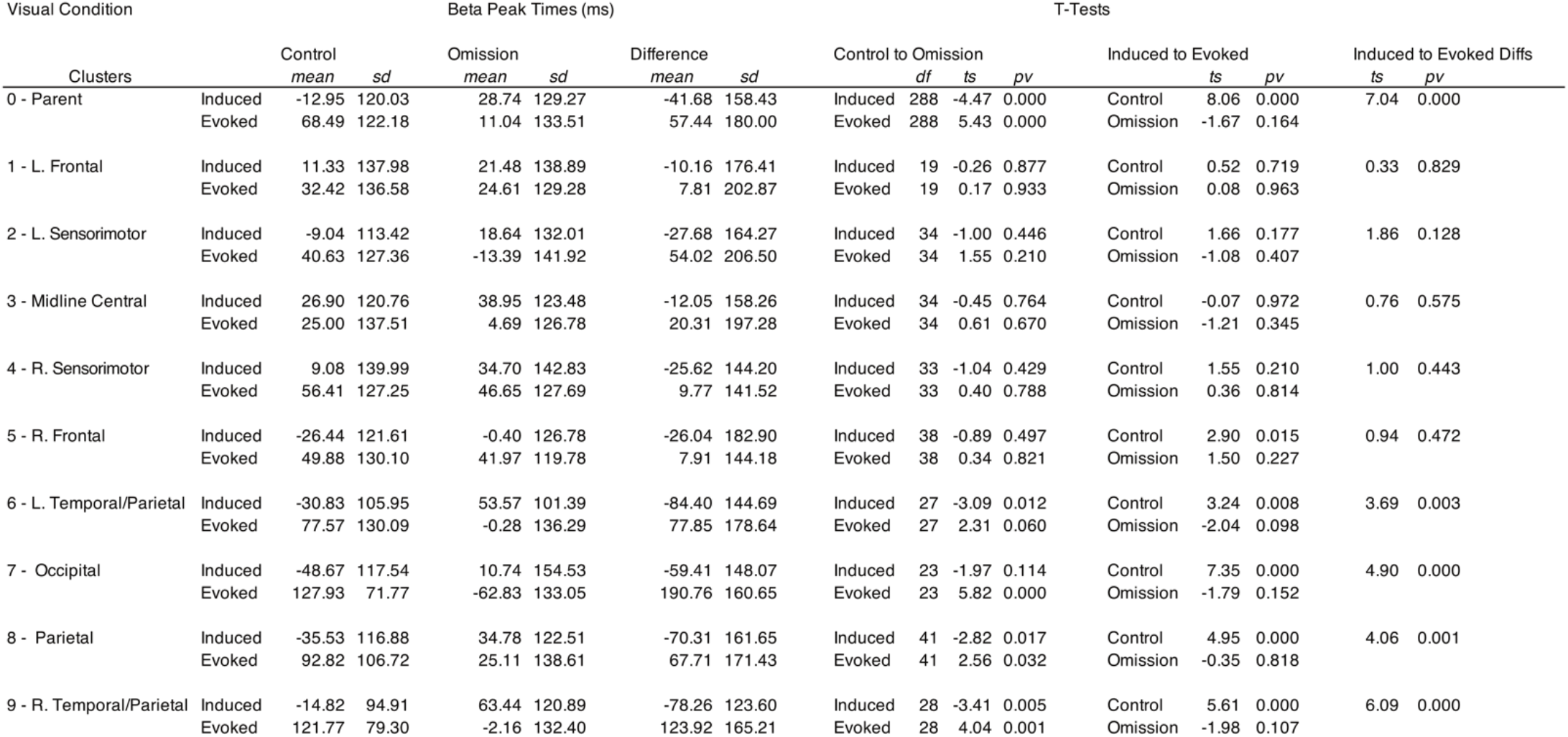
Visual peak times for beta activity within +/− 200 ms of event or omission onset. Difference values are calculated as the difference between control and omission times. Control to omission tests comparisons are made for both induced and evoked activity. Induced to evoked test comparisons are made for both control and omission conditions. Induced to evoke difference tests are made between the difference values calculated. Degrees of freedom were the same for each of the t-tests on the same row. P-values displayed are FDR corrected.

**Supplemental Table 4.**
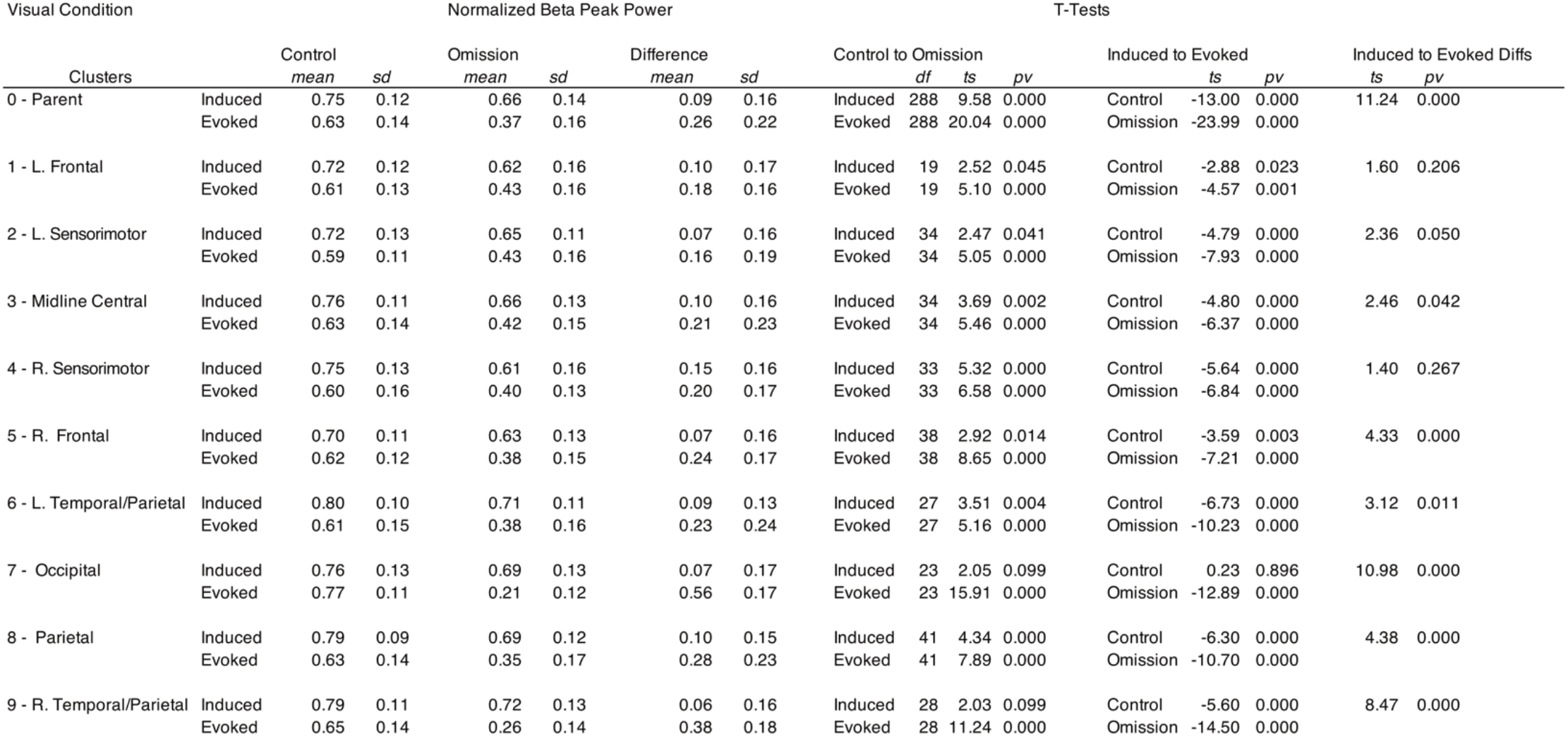
Normalized visual peak times for beta activity within +/− 200 ms of event or omission onset. Difference values are calculated as the difference between control and omission times. Control to omission tests comparisons are made for both induced and evoked activity. Induced to evoked test comparisons are made for both control and omission conditions. Induced to evoke difference tests are made between the difference values calculated. Degrees of freedom were the same for each of the t-tests on the same row. P-values displayed are FDR corrected.

**Supplemental Table 5.**
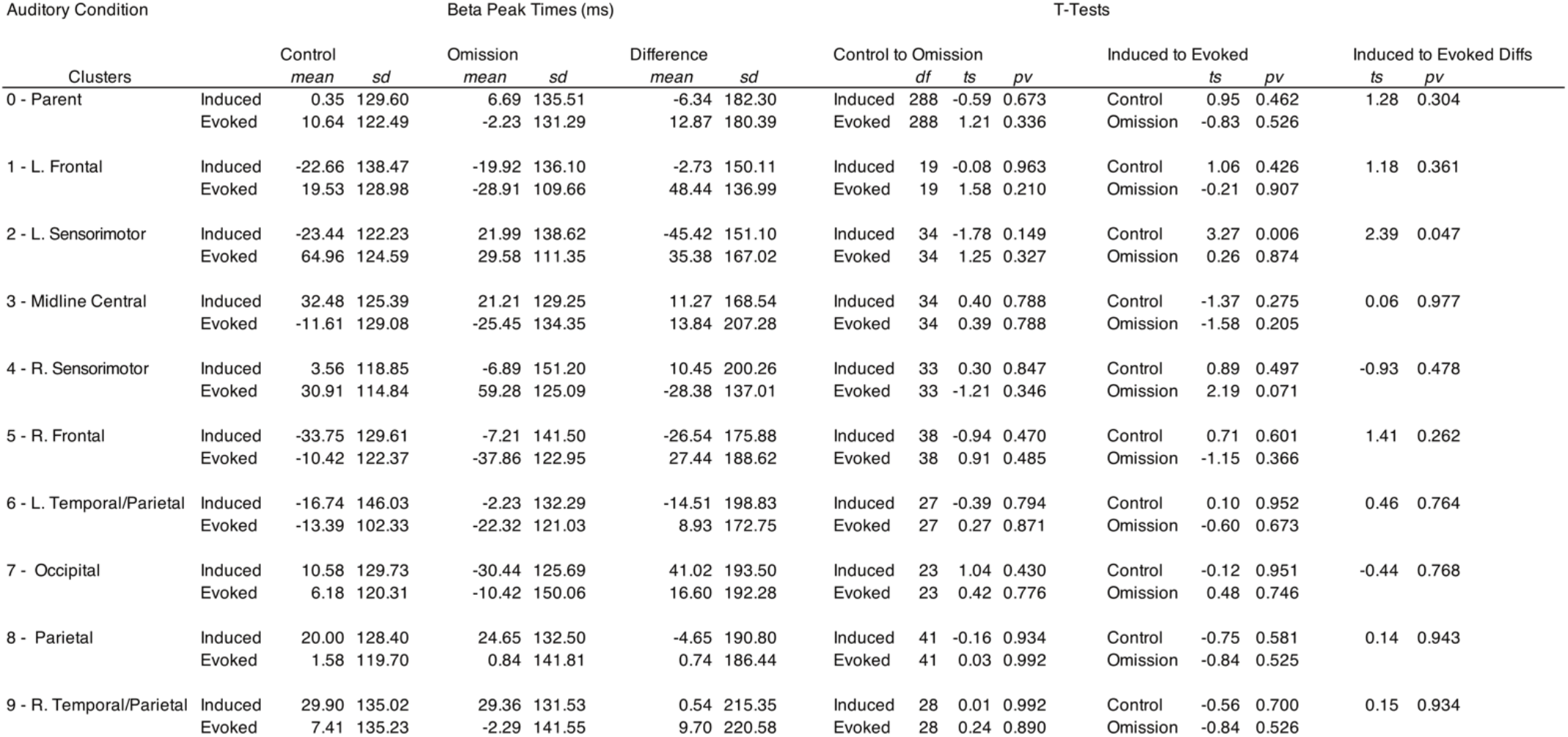
Auditory peak times for beta activity within +/− 200 ms of event or omission onset. Difference values are calculated as the difference between control and omission times. Control to omission tests comparisons are made for both induced and evoked activity. Induced to evoked test comparisons are made for both control and omission conditions. Induced to evoke difference tests are made between the difference values calculated. Degrees of freedom were the same for each of the t-tests on the same row. P-values displayed are FDR corrected.

**Supplemental Table 6.**
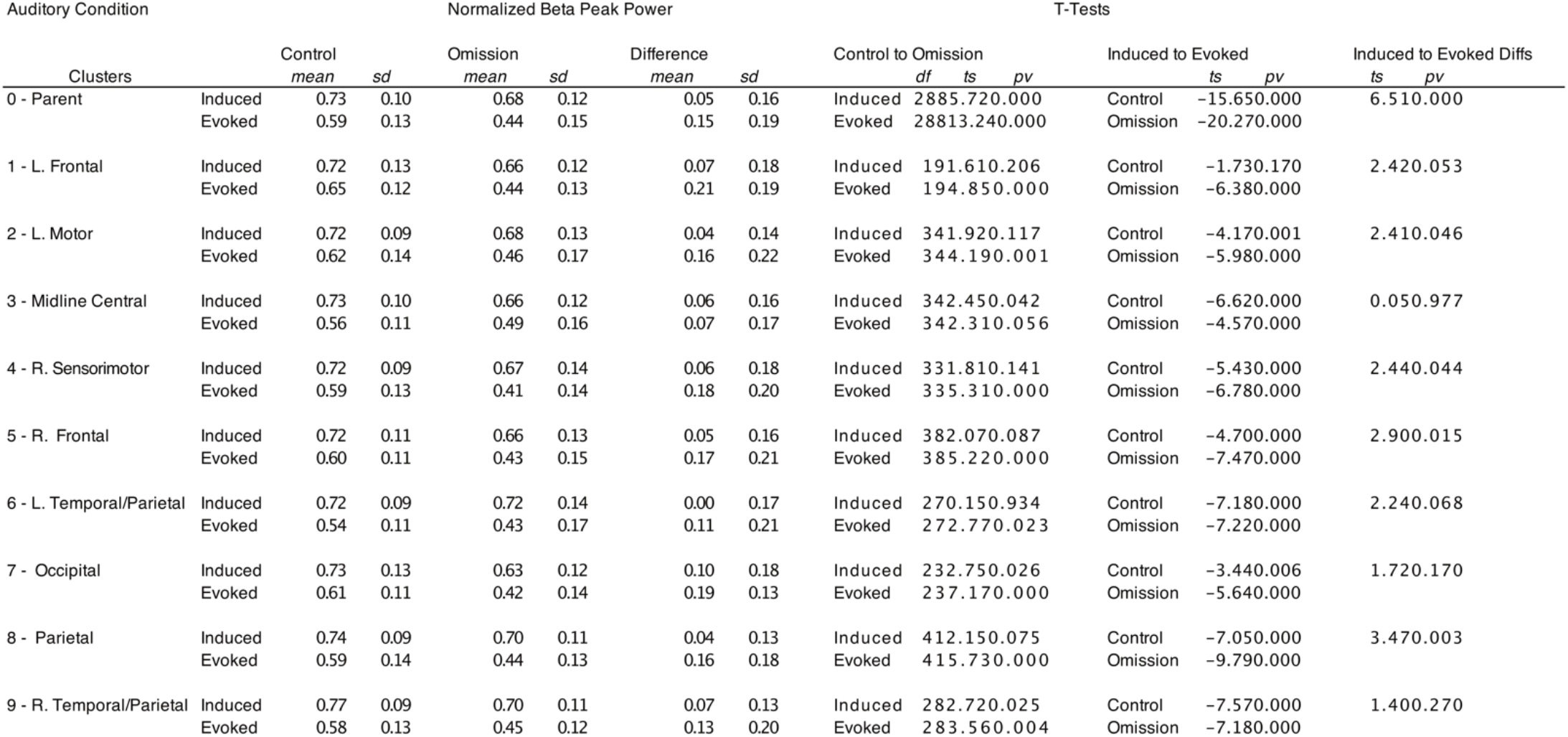
Normalized auditory peak times for beta activity within +/− 200 ms of event or omission onset. Difference values are calculated as the difference between control and omission times. Control to omission tests comparisons are made for both induced and evoked activity. Induced to evoked test comparisons are made for both control and omission conditions. Induced to evoke difference tests are made between the difference values calculated. Degrees of freedom were the same for each of the t-tests on the same row. P-values displayed are FDR corrected

## References

Abbott, N. T., & Shahin, A. J. (2018). Cross-modal phonetic encoding facilitates the McGurk illusion and phonemic restoration. Journal of neurophysiology, 120(6), 2988–3000.

Araneda, R., Renier, L., Ebner-Karestinos, D., Dricot, L., and De Volder, A. G. (2017). Hearing, feeling or seeing a beat recruits a supramodal network in the auditory dorsal stream. Eur. J. Neurosci. 45, 1439–1450.

Arnal, L. H., Doelling, K. B., & Poeppel, D. (2014). Delta–beta coupled oscillations underlie temporal prediction accuracy. Cerebral Cortex, 25(9), 3077–3085.

Arnal, L. H., & Giraud, A. L. (2012). Cortical oscillations and sensory predictions. Trends in cognitive sciences, 16(7), 390–398.

Arnal, L. H., Wyart, V., & Giraud, A. L. (2011). Transitions in neural oscillations reflect prediction errors generated in audiovisual speech. Nature neuroscience, 14(6), 797.

Bastos, A. M., Vezoli, J., Bosman, C. A., Schoffelen, J. M., Oostenveld, R., Dowdall, J. R., … & Fries, P. (2015). Visual areas exert feedforward and feedback influences through distinct frequency channels. Neuron, 85(2), 390–401.

Bauer, A. K. R., Bleichner, M. G., Jaeger, M., Thorne, J. D., & Debener, S. (2018). Dynamic phase alignment of ongoing auditory cortex oscillations. Neuroimage, 167, 396–407.

Benjamini, Y., & Hochberg, Y. (1995). Controlling the false discovery rate: a practical and powerful approach to multiple testing. Journal of the Royal statistical society: series B (Methodological), 57(1), 289–300.

Chang, C. Y., Hsu, S. H., Pion-Tonachini, L., & Jung, T. P. (2018, July). Evaluation of artifact subspace reconstruction for automatic EEG artifact removal. In 2018 40th Annual International Conference of the IEEE Engineering in Medicine and Biology Society (EMBC) (pp. 1242–1245). IEEE.

Chen, Y., Repp, B. H., and Patel, A. D. (2002). Spectral decomposition of variability in synchronization and continuation tapping: comparisons between auditory and visual pacing and feedback conditions. Hum. Mov. Sci. 21, 515–532.

Comstock, D. C., and Balasubramaniam, R. (2018). Neural responses to perturbations in visual and auditory metronomes during sensorimotor synchronization. Neuropsychologia 117, 55–66.

Comstock, D. C., Hove, M. J., & Balasubramaniam, R. (2018). Sensorimotor synchronization with auditory and visual modalities: Behavioral and neural differences. Frontiers in computational neuroscience, 12.

Delorme, A., & Makeig, S. (2004). EEGLAB: an open source toolbox for analysis of single-trial EEG dynamics including independent component analysis. Journal of neuroscience methods, 134(1), 9–21.

Destoky, F., Philippe, M., Bertels, J., Verhasselt, M., Coquelet, N., Vander Ghinst, M., … & Bourguignon, M. (2019). Comparing the potential of MEG and EEG to uncover brain tracking of speech temporal envelope. Neuroimage, 184, 201–213.

Engel, A. K., & Fries, P. (2010). Beta-band oscillations—signalling the status quo?. Current opinion in neurobiology, 20(2), 156–165.

Fitch, W. (2016). Dance, music, meter and groove: a forgotten partnership. Frontiers in human neuroscience, 10, 64.

Fujioka, T., Ross, B., & Trainor, L. J. (2015). Beta-band oscillations represent auditory beat and its metrical hierarchy in perception and imagery. Journal of Neuroscience, 35(45), 15187–15198.

Fujioka, T., Trainor, L., Large, E., & Ross, B. (2009). Beta and gamma rhythms in human auditory cortex during musical beat processing. Annals of the New York Academy of Sciences, 1169(1), 89–92.

Fujioka, T., Trainor, L. J., Large, E. W., & Ross, B. (2012). Internalized timing of isochronous sounds is represented in neuromagnetic beta oscillations. Journal of Neuroscience, 32(5), 1791–1802.

Gan, L., Huang, Y., Zhou, L., Qian, C., & Wu, X. (2015). Synchronization to a bouncing ball with a realistic motion trajectory. Scientific reports, 5, 11974.

Gelman, A., & Stern, H. (2006). The difference between “significant” and “not significant” is not itself statistically significant. The American Statistician, 60(4), 328–331.

Gordon, C. L., Cobb, P. R., & Balasubramaniam, R. (2018). Recruitment of the motor system during music listening: An ALE meta-analysis of fMRI data. PloS one, 13(11), e0207213.

Grube, M., Cooper, F. E., Chinnery, P. F., & Griffiths, T. D. (2010). Dissociation of duration-based and beat-based auditory timing in cerebellar degeneration. Proceedings of the National Academy of Sciences, 107(25), 11597–11601.

Grube, M., Lee, K. H., Griffiths, T. D., Barker, A. T., & Woodruff, P. W. (2010). Transcranial magnetic theta-burst stimulation of the human cerebellum distinguishes absolute, duration-based from relative, beat-based perception of subsecond time intervals. Frontiers in Psychology, 1, 171.

Gruzelier, J. H., Holmes, P., Hirst, L., Bulpin, K., Rahman, S., Van Run, C., & Leach, J. (2014). Replication of elite music performance enhancement following alpha/theta neurofeedback and application to novice performance and improvisation with SMR benefits. Biological psychology, 95, 96–107.

Haegens, S., & Golumbic, E. Z. (2018). Rhythmic facilitation of sensory processing: a critical review. Neuroscience & Biobehavioral Reviews, 86, 150–165.

Hove, M. J., Fairhurst, M. T., Kotz, S. A., and Keller, P. E. (2013a). Synchronizing with auditory and visual rhythms: an fMRI assessment of modality differences and modality appropriateness. Neuroimage 67, 313–321.

Hove, M. J., Iversen, J. R., Zhang, A., and Repp, B. H. (2013b). Synchronization with competing visual and auditory rhythms: bouncing ball meets metronome. Psychol. Res. 77, 388–398.

Hove, M. J., Spivey, M. J., & Krumhansl, C. L. (2010). Compatibility of motion facilitates visuomotor synchronization. Journal of Experimental Psychology: Human Perception and Performance, 36(6), 1525.

Iversen, J. R. (2016). In the beginning was the beat: evolutionary origins of musical rhythm in humans. in Hartenberger, R. The Cambridge Companion to Percussion, Cambridge University Press, Cambridge, 281–295.

Iversen, J. R., & Balasubramaniam, R. (2016). Synchronization and temporal processing. Current Opinion in Behavioral Sciences, 8, 175–180.

Iversen, J. R., Patel, A. D., Nicodemus, B., and Emmorey, K. (2015). Synchronization to auditory and visual rhythms in hearing and deaf individuals. Cognition 134, 232–244.

Iversen, J., Repp, B., & Patel, A. (2009). Top-down control of rhythm perception modulates early auditory responses. Annals of the New York Academy of Sciences, 1169(1), 58–73.

Janata, P., Tomic, S. T., & Haberman, J. M. (2012). Sensorimotor coupling in music and the psychology of the groove. Journal of Experimental Psychology: General, 141(1), 54.

Jäncke, L., Loose, R., Lutz, K., Specht, K., & Shah, N. J. (2000). Cortical activations during paced finger-tapping applying visual and auditory pacing stimuli. Cognitive Brain Research, 10(1-2), 51–66.

Jantzen, K. J., Steinberg, F. L., & Kelso, J. A. S. (2005). Functional MRI reveals the existence of modality and coordination-dependent timing networks. Neuroimage, 25(4), 1031–1042.

Kilavik, B. E., Zaepffel, M., Brovelli, A., MacKay, W. A., & Riehle, A. (2013). The ups and downs of beta oscillations in sensorimotor cortex. Experimental neurology, 245, 15–26.

Kopell, N., Ermentrout, G. B., Whittington, M. A., & Traub, R. D. (2000). Gamma rhythms and beta rhythms have different synchronization properties. Proceedings of the National Academy of Sciences, 97(4), 1867–1872.

D.J. Levitin, J.A. Grahn, J. London. The psychology of music: Rhythm and movement. (2018) Annual Review of Psychology, 69:51–75.

Lorås, H., Sigmundsson, H., Talcott, J. B., Öhberg, F., & Stensdotter, A. K. (2012). Timing continuous or discontinuous movements across effectors specified by different pacing modalities and intervals. Experimental brain research, 220(3-4), 335–347.

Meijer, D., Te Woerd, E., & Praamstra, P. (2016). Timing of beta oscillatory synchronization and temporal prediction of upcoming stimuli. NeuroImage, 138, 233–241.

Merchant H, Perez O, Zarco W, Gamez J. 2013 Interval tuning in the primate medial premotor cortex as a general timing mechanism. J. Neurosci. 33, 9082–9096.

Merchant, H., Grahn, J., Trainor, L., Rohrmeier, M., & Fitch, W. T. (2015). Finding the beat: A neural perspective across humans and non-human primates. Philosophical Transactions of the Royal Society B: Biological Sciences, 370(1664).

Michalareas, G., Vezoli, J., Van Pelt, S., Schoffelen, J. M., Kennedy, H., & Fries, P. (2016). Alpha-beta and gamma rhythms subserve feedback and feedforward influences among human visual cortical areas. Neuron, 89(2), 384–397.

Morillon, B., Hackett, T. A., Kajikawa, Y., & Schroeder, C. E. (2015). Predictive motor control of sensory dynamics in auditory active sensing. Current Opinion in Neurobiology, 31, 230–238.

Mullen, T. R., Kothe, C. A., Chi, Y. M., Ojeda, A., Kerth, T., Makeig, S., … & Cauwenberghs, G. (2015). Real-time neuroimaging and cognitive monitoring using wearable dry EEG. IEEE Transactions on Biomedical Engineering, 62(11), 2553–2567.

Ng, B. S. W., Schroeder, T., & Kayser, C. (2012). A precluding but not ensuring role of entrained low-frequency oscillations for auditory perception. Journal of Neuroscience, 32(35), 12268–12276.

Obleser, J., Eisner, F., & Kotz, S. A. (2008). Bilateral speech comprehension reflects differential sensitivity to spectral and temporal features. Journal of neuroscience, 28(32), 8116–8123.

Palmer, J. A., Kreutz-Delgado, K., & Makeig, S. (2012). AMICA: An adaptive mixture of independent component analyzers with shared components. Swartz Center for Computatonal Neursoscience, University of California San Diego, Tech. Rep.

Patel, A. D. (2006). Musical rhythm, linguistic rhythm, and human evolution. Music Perception: An Interdisciplinary Journal, 24(1), 99–104.

Patel, A. D., & Iversen, J. R. (2014). The evolutionary neuroscience of musical beat perception: the Action Simulation for Auditory Prediction (ASAP) hypothesis. Frontiers in systems neuroscience, 8, 57.

Perception Research Systems. 2007. Paradigm Stimulus Presentation, Retrieved from http://www.paradigmexperiments.com

Piazza, C., Miyakoshi, M., Akalin-Acar, Z., Cantiani, C., Reni, G., Bianchi, A. M., & Makeig, S. (2016). An automated function for identifying eeg independent components representing bilateral source activity. In XIV Mediterranean Conference on Medical and Biological Engineering and Computing 2016 (pp. 105–109). Springer, Cham.

Picazio, S., Veniero, D., Ponzo, V., Caltagirone, C., Gross, J., Thut, G., & Koch, G. (2014). Prefrontal control over motor cortex cycles at beta frequency during movement inhibition. Current Biology, 24(24), 2940–2945.

Repp, B. H. (2003). Rate limits in sensorimotor synchronization with auditory and visual sequences: the synchronization threshold and the benefits and costs of interval subdivision. J. Motor Behav. 35, 355–370.

Repp, B. H., and Penel, A. (2004). Rhythmic movement is attracted more strongly to auditory than to visual rhythms. Psychol. Res. 68, 252–270.

Repp, B. H., and Su, Y. H. (2013). Sensorimotor synchronization: a review of recent research (2006-2012). Psychon. Bull. Rev. 20, 403–452

Riecke, L., Sack, A. T., & Schroeder, C. E. (2015). Endogenous delta/theta sound-brain phase entrainment accelerates the buildup of auditory streaming. Current Biology, 25(24), 3196–3201.

Ross, J. M., Iversen, J. R., and Balasubramaniam, R. (2016a). Motor simulation theories of musical beat perception. Neurocase 22, 558–565.

Ross, J. M., Iversen, J. R., and Balasubramaniam, R. (2018). The role of posterior parietal cortex in beat-based timing perception: a continuous theta burst stimulation study. J. Cogn. Neurosci. 30, 634–643.

Ross, J. M., Warlaumont, A. S., Abney, D. H., Rigoli, L. M., & Balasubramaniam, R. (2016). Influence of musical groove on postural sway. Journal of Experimental Psychology: Human Perception and Performance, 42(3), 308.

Saleh, M., Reimer, J., Penn, R., Ojakangas, C. L., & Hatsopoulos, N. G. (2010). Fast and slow oscillations in human primary motor cortex predict oncoming behaviorally relevant cues. Neuron, 65(4), 461–471.

Sarnthein, J., Petsche, H., Rappelsberger, P., Shaw, G. L., & Von Stein, A. (1998). Synchronization between prefrontal and posterior association cortex during human working memory. Proceedings of the National Academy of Sciences, 95(12), 7092–7096.

Schroeder, C. E., & Lakatos, P. (2009). Low-frequency neuronal oscillations as instruments of sensory selection. Trends in neurosciences, 32(1), 9–18.

Schubotz, R. I., Friederici, A. D., & Von Cramon, D. Y. (2000). Time perception and motor timing: a common cortical and subcortical basis revealed by fMRI. Neuroimage, 11(1), 1–12.

Snyder, J. S., & Large, E. W. (2005). Gamma-band activity reflects the metric structure of rhythmic tone sequences. Cognitive brain research, 24(1), 117–126.

Tomassini, A., Ambrogioni, L., Medendorp, W. P., & Maris, E. (2017). Theta oscillations locked to intended actions rhythmically modulate perception. Elife, 6, e25618.

Varlet, M., Nozaradan, S., Trainor, L., & Keller, P. E. (2020). Dynamic Modulation of Beta Band Cortico-Muscular Coupling Induced by Audio–Visual Rhythms. Cerebral Cortex Communications, 1(1), tgaa043.

von Stein, A., and Sarnthein, J. (2000). Different frequencies for different scales of cortical integration: from local gamma to long range alpha/theta synchronization. Int. J. Psychophysiol. 38, 301–313.

Wiener, M., and Kanai, R. (2016). Frequency tuning for temporal perception and prediction. Curr. Opin. Behav. Sci. 8, 1–6.

Wiener, M., Turkeltaub, P., & Coslett, H. B. (2010). The image of time: a voxel-wise meta-analysis. Neuroimage, 49(2), 1728–1740.

Zatorre, R. J., & Belin, P. (2001). Spectral and temporal processing in human auditory cortex. Cerebral cortex, 11(10), 946–953.

Zatorre, R. J., Evans, A. C., Meyer, E., & Gjedde, A. (1992). Lateralization of phonetic and pitch discrimination in speech processing. Science, 256(5058), 846–849.

Zhou, B., Yang, S., Mao, L., & Han, S. (2014). Visual feature processing in the early visual cortex affects duration perception. Journal of Experimental Psychology: General, 143(5), 1893.

